# Integrative taxonomy of the North American pink flat-backed millipedes (Polydesmidae: *Pseudopolydesmus* Attems, 1898)

**DOI:** 10.64898/2026.10.30.763145

**Authors:** Derek A. Hennen, Paul E. Marek

**Author notes:** Virginia Department of Conservation and Recreation, Natural Heritage Program, Richmond, Virginia 23219.

## Abstract

Millipedes of the genus *Pseudopolydesmus* Attems, 1898 are common millipedes in North America. They are the largest, in terms of body size, and most geographically wide-ranging of the family Polydesmidae, and are ubiquitous in eastern and central North American forests. Eight species have been named based on the morphology of the genitalia, but these have not been investigated genetically. Here, we take an integrated taxonomic approach, using five genes to infer a molecular phylogeny of *Pseudopolydesmus* species, and analyze a dataset of 15 morphological characters in an effort to delimit species in the genus. 1,148 *Pseudopolydesmus* specimens were newly collected for this study, and all known species were sampled. These specimens were used for analysis, and were combined with museum specimens and literature records into a database of 3,462 unique specimen records. This database was used to produce distribution maps for each species in the genus. The maximum likelihood phylogeny found support for the eight species in the genus, supporting previous morphological species hypotheses. However, previous morphology-based hypotheses regarding evolutionary relationships between the species were not supported by the analysis. The morphological analysis, performed with a modified PCA, did not delimit species clusters as well as the genitalic and genetic differences did, due to comparatively uniform somatic morphology in the genus. However, our results show that genitalic morphology, combined with genetic analysis, provide congruent species delimitation hypotheses for *Pseudopolydesmus*, and these lines of evidence may prove useful in delimiting species and generic boundaries in other Polydesmidae worldwide.

## Introduction

The millipede genus *Pseudopolydesmus* (Attems, 1898) is one of the most commonly-encountered millipedes in central and eastern North America. With eight nominative species, the genus covers a large geographic area (Fig. 1) from North Dakota south through Texas, east to Florida, and north to Québec. They are typically pink or brown in color, but one species, *Pseudopolydesmus canadensis* (Newport, 1844), often has deep red paranota (Fig. 2). *Pseudopolydesmus* species range between 7 – 32 mm in length, and are found under logs and in leaf litter in deciduous forests (Withrow 1988).

**Figure 1.**
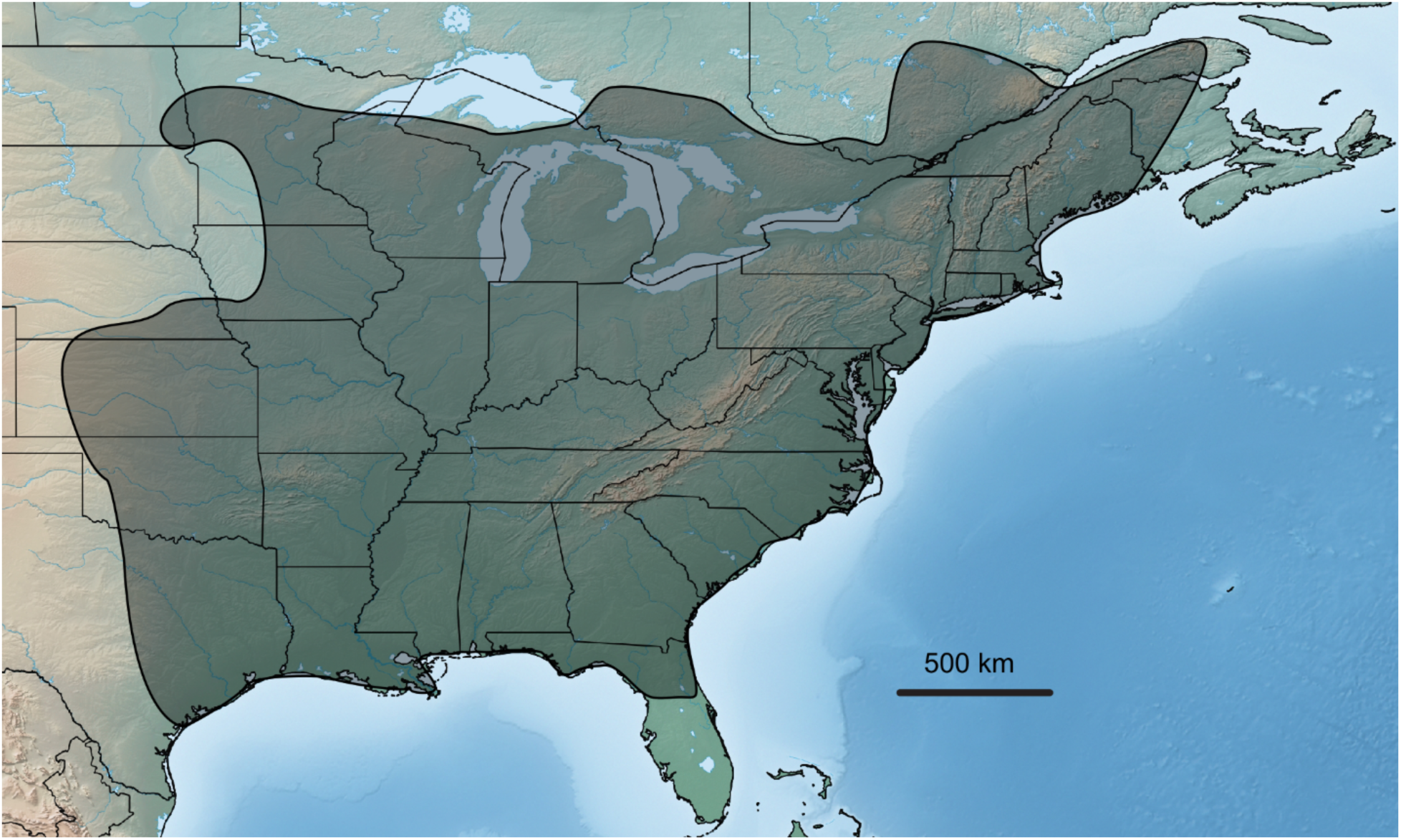
Distribution of the genus *Pseudopolydesmus* in eastern North America, shaded black.

**Figure 2.**
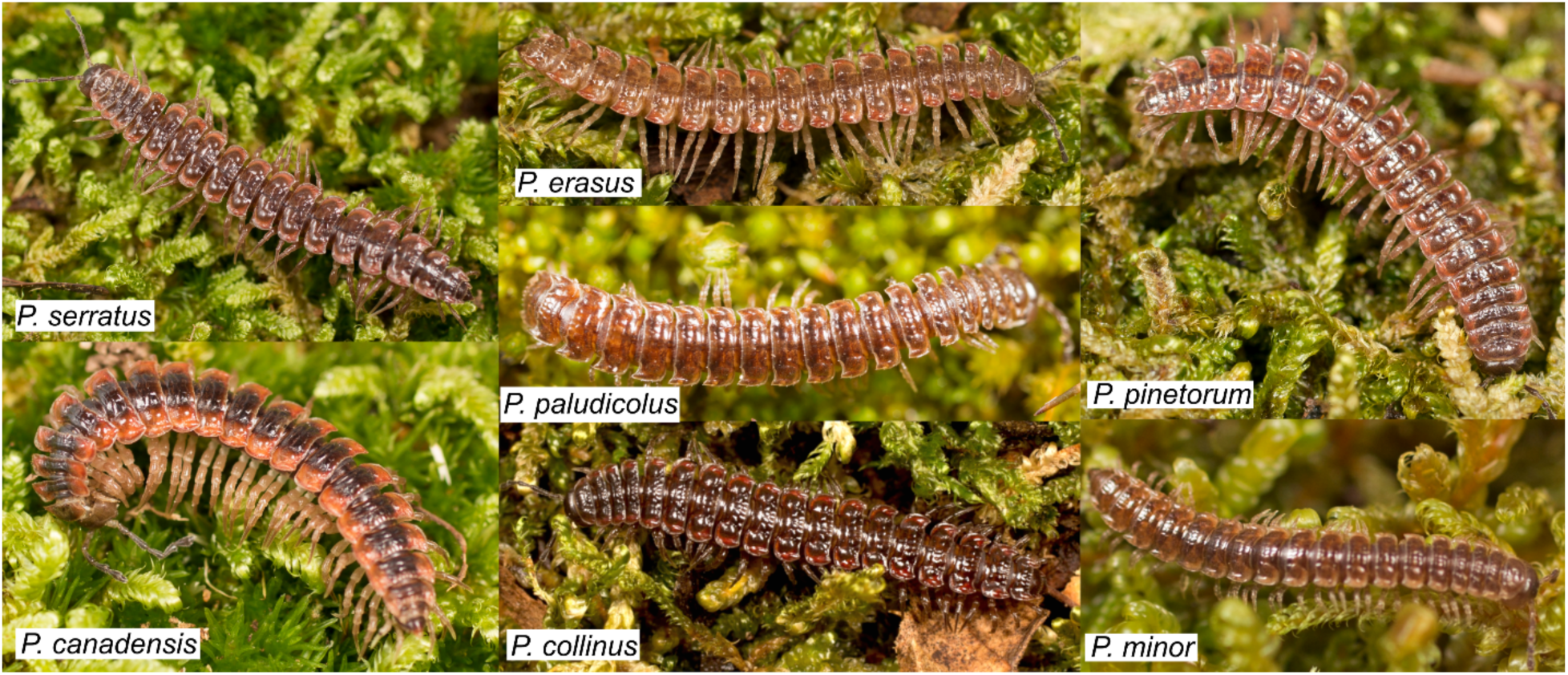
General habitus of *Pseudopolydesmus* species, not to scale. *Pseudopolydesmus caddo* not shown.

A historical review of the genus, synthesizing available taxonomic information up to 2019, has been provided by Sierwald et al. (2019); this provided a key and lists of synonymies for each species, photographs and scanning electron micrographs of the gonopods of each species, and comments on morphological terminology in the genus. Hence, only a brief summary of the taxonomic history of the genus is given here. Eight species of *Pseudopolydesmus* are currently recognized, based purely on morphological characters: *Pseudopolydesmus serratus* (Say, 1821), *Pseudopolydesmus canadensis, Pseudopolydesmus minor* (Bollman, 1888), *Pseudopolydesmus pinetorum* (Bollman, 1888), *Pseudopolydesmus erasus* (Loomis, 1943), *Pseudopolydesmus caddo* Chamberlin, 1949, *Pseudopolydesmus paludicolus* Hoffman, 1950, and *Pseudopolydesmus collinus* Hoffman, 1974. The first species to be described, *P. serratus*, was placed in the genus *Polydesmus* Latreille, 1802, and was collected on the eastern shore of Virginia (Sierwald et al. 2019). Most names in the genus were described under the genera *Polydesmus* or *Dixidesmus* Chamberlin, 1943 (the latter now recognized as a junior subjective synonym of *Pseudopolydesmus*), and 32 species names have been ascribed to *Pseudopolydesmus*, with 18 of those names coined by R.V. Chamberlin. Most work on *Pseudopolydesmus* consisted of short taxonomic descriptions of a single species or a few species. Hoffman (1974) established a useful terminology for the gonopod processes in his description of *P. collinus*, which was subsequently adopted by taxonomists working on the genus. An unpublished Ph.D. dissertation by Withrow (1988) summarized the distribution of the genus, was the first work to provide a key to species, provided ecological information, and hypothesized relationships within the genus and with other Polydesmidae. However, it was never published, limiting its usefulness, and preventing its changes to the classification.

Sporadic interest in the genus during the early 21th century provided useful information on its natural history from laboratory observations (Youngsteadt 2009), and a litany of piecemeal distributional records from across its range (Shelley & Snyder 2012). The review by Sierwald et al. (2019) postulated species within the genus and provided clear morphological images for species identification. This resolved the confusion that has continually trailed the genus since the description of its first species. Most recently, the genital morphology of female *Pseudopolydesmus* millipedes and the hypothesized mechanics of mating in the genus were investigated by Zahnle et al. (2020). This investigation augmented our understanding of a poorly understood area of study in millipedes.

Despite these advances, phylogenetic relationships within the genus *Pseudopolydesmus* and the family Polydesmidae remain understudied. The family is mainly Holarctic, with about 30 described genera and over 240 species (Enghoff et al. 2015). The group is not well-delimited taxonomically, never has been analyzed with molecular phylogenetics, and has been described as a wastebasket (Shear 2012) due to large species-rich genera such as *Polydesmus* and *Epanerchodus* Attems, 1901; these genera have many ill-defined species and subgenera. A subfamilial classification was proposed by Withrow (1988) and referenced by later workers (Simonsen 1990; Golovatch 1991), but was not officially published, and has not been widely adopted (Shelley 1996a). Much synthetic work remains to be done—preferably supported by molecular phylogenetics—particularly with the European and Asian genera. While historically challenging, the genera of Polydesmidae present an excellent opportunity for using molecular phylogenetics to bring order to the historical chaos of the genus.

In North America, the taxonomy of the family has received more recent work, with 10 precinctive genera: *Pseudopolydesmus*; *Scytonotus* Koch, 1847; *Bidentogon* Buckett & Gardner, 1968; *Calianotus* Shelley, 1997; *Rowlandesmus* Shear & Marek, 2021; *Retrorsia* Shelley, 2003; *Retrorsioides* Shear & Marek, 2021; *Benedictesmus* Shear & Marek, 2021; *Snoqualmia* Shear, 2012; and *Utadesmus* Chamberlin & Hoffman, 1950. There are two introduced genera: *Polydesmus* and *Brachydesmus* Heller, 1858. Shear (2012) and Shear & Reddell (2017) provided keys to the genera. The native North American genera are comparably well-delineated and have clear illustrations of their morphology, allowing for confident identification (Chamberlin 1930; Chamberlin & Hoffman 1950; Buckett & Gardner 1968; Shelley 1993, 1997, 2003; Shear 2012; Shear & Marek, 2021). Little is known about the ecology of these genera, and most information on them is taxonomic and distributional in nature, but *Scytonotus* has been revised (Shelley 1993); this genus is the only native polydesmid genus found in both western and eastern North America. New genera and species from western North America await description (Shear & Reddell 2017), presenting another opportunity for the use of molecular phylogenetics to clarify intergeneric relationships.

Within *Pseudopolydesmus*, hypothesized relationships between species have been based on morphological characters, especially those of the gonopods (Withrow 1988, Sierwald et al. 2019). Two main species groups have been proposed: the *canadensis* group and the *serratus* group. The *canadensis* group is composed of *Pseudopolydesmus canadensis, P. collinus, P. erasus*, and *P. pinetorum*, while the *serratus* group contains *Pseudopolydesmus serratus, P. caddo, P. minor,* and *P. paludicolus*. Withrow summarized his hypothesized relationships as follows: ((*pinetorum*, ((*tallulanus*, *erasus*), (*canadensis*, *collinus*))), ((*serratus*, *paludicolus*, (*minor*, *caddo*)) (1988, his figure 121). *Pseudopolydesmus tallulanus* (Chamberlin, 1943) was determined to be a junior subjective synonym of *P. erasus* by Sierwald et al. (2019). These relationships have not been tested by molecular phylogenetics, and remain the sole hypothesis for species delimitation within the genus.

Species delimitation in millipedes has historically placed emphasis on gonopod morphology, but genetic investigations have revealed cryptic species complexes in juliformian taxa with uniform gonopod morphology (Bond & Sierwald 2002, Walker et al. 2009, Mwabvu et al. 2013, 2015). Within the Polydesmidae, a proliferation of inconsistently-applied terms for the various processes of the gonopods has emerged, causing difficulties in direct comparison of species. As a result, homologies of the gonopod structures are uncertain, and challenging to efficiently compare, despite efforts to standardize the terminology (see Shear & Marek, 2021).

The polydesmid gonopod ranges from a relatively simple, singly-branched structure with a few triangular or quadrate processes (such as in *Pseudopolydesmus* or *Pacidesmus* Golovatch, 1991) to a complex, multi-branched structure with many differently-shaped processes (such as in *Snoqualmia*). A few terms, such as coxa and cannula, have been consistently applied throughout the family. Other terms, such as endomerite and acropodite, are not as consistently applied. The acropodite, for example, has also been called the tibiotarsus (Shelley, 2003), endomere (Anh, 2009), and distofemoral part (Djursvoll, 2008). Thus, the current suite of descriptive terms is large and open to interpretation, and species delimitation and description within the Polydesmidae (and within the suborder Polydesmidea) remains subjective.

The use of objective and repeatable methods for species delimitation provides a path for solving this problem. Recent studies have highlighted the usefulness of genetic data for species delimitation and identification, with the cytochrome c oxidase subunit I (COI) gene proving useful for identifying polydesmid species (Spelda et al. 2011; Angyal et al. 2017). A modified principal components analysis (PCA) for species delimitation based on morphological measurements has recently been developed (Ezard et al. 2010) and used in an integrative framework with genetic data to revise the Opiliones genus *Sclerobunus* in western North America (Derkarabetian & Hedin 2014). Like millipedes, Opiliones are low-vagility organisms, not able to fly or disperse widely, and with many species distributions closely corresponding to the physiography of their region.

In this study, our aim is to delimit species of *Pseudopolydesmus* in a framework integrating molecular phylogenetics and morphology. Here, we test the monophyly of *Pseudopolydesmus*, provide distributions for each species, photographs of living individuals, and ecological information for each species. Useful morphological characters for delimiting species in the genus, and possible characters for use throughout the family are also discussed.

## Materials and Methods

Specimens of *Pseudopolydesmus* were collected in 20 states in the central and eastern United States from 2013 – 2019, according to methods outlined in Means et al. (2015). Mesic deciduous forests were the primary habitats targeted for field collection, and specimens were searched for under leaf litter and logs. Each putative species in the genus was targeted for recollecting, and historical holotype localities were visited when possible. Old notes from Richard Hoffman were examined for information on any possible new species in the genus, and one potential new species was noted by Hoffman from southwest Virginia. This possible new species was also targeted during field collecting. Specimens were collected throughout the range of each species in order to obtain ample specimens for population analyses.

After field collection, specimens were kept alive for processing in the lab. When living specimens were returned to the lab, representative specimens were photographed with a Canon 6D dSLR camera with a 50 mm or 65 mm Canon MP-E macro lens. The 50 mm lens was used for medium to large-sized individuals, while the 65 mm lens was used for small individuals.

Specimens were subsequently given a catalog number beginning with the prefix MPE, and 24 legs were removed from the right side of the body and preserved in 100% ethanol or RNAlater (Qiagen) for molecular analysis, while the rest of the body was preserved in 70% isopropanol for morphological analysis and deposited in the Virginia Tech Insect Collection (abbreviation VTEC, https://www.collection.ento.vt.edu).

In addition to field-collected specimens, museum material and literature records were also examined. Museum specimens were borrowed from the Virginia Museum of Natural History (VMNH, Martinsville, Virginia), the North Carolina Museum of Natural Sciences (NCSM, Raleigh, North Carolina), and the Illinois Natural History Survey (INHS, Champaign, Illinois). Specimens from the VMNH and NCSM were georeferenced with the program GEOLocate (http://www.museum.tulane.edu/geolocate/web/WebGeoref.aspx), and databased with catalog numbers beginning with the prefix PSE (e.g. PSE0001). Additional specimen records were downloaded from the Symbiota Collections of Arthropods Network (SCAN) database (https://scan-bugs.org/). Literature records from published works on *Pseudopolydesmus* (Shelley & Snyder 2012, Withrow 1988) were digitized, georeferenced as above, and included with museum records into a distributional database of the genus. Distribution maps for each species were produced with the website SimpleMappr (Shorthouse, 2010).

Field-collected specimens were processed for molecular phylogenetics according to previously published methods (Means & Marek, 2017; Means et al., 2021). Outgroup taxa were selected from within the Polydesmidae and from additional families in the suborder Polydesmidea. Six genera from the family Polydesmidae were included: ten species of *Epanerchodus*, 3 species of *Polydesmus*, 1 species of *Brachydesmus*, 2 species of *Schizoturanius* Verhoeff, 1931, 1 species of *Scytonotus*, and 1 unknown genus and species of Polydesmidae.

Five families from the suborder Polydesmidea were included: 4 species of Macrosternodesmidae, 2 species of Opisotretidae, 2 species of Cryptodesmidae, 2 species of Haplodesmidae, and 1 species of Pyrgodesmidae. An unidentified species from the suborder Dalodesmidea, family Dalodesmidae was used to root the phylogeny. Genomic DNA from 4-6 legs was extracted and purified with a Qiagen DNeasy kit, according to instructions of the manufacturer. For very small specimens, genomic DNA was extracted from 4-8 body rings after removing the gut. DNA was amplified with polymerase chain reaction, and bands visualized with an agarose gel to confirm product amplification. Amplified DNA was cleaned, concentrations quantified and normalized, and then sequenced with an Applied Biosystems 3730 DNA Analyzer at the University of Arizona Genetics Core. Five gene regions were sequenced: two mitochondrial genes, cytochrome c oxidase subunit I (COI) and large subunit ribosomal RNA (16S); and three nuclear genes: 28S ribosomal RNA (28S), elongation factor-1 alpha (EF1-a), and Enolase, with amplification protocols found in Means & Marek (2017). Enolase was chosen due to its successful use in Coleoptera phylogenetics (Wild & Maddison, 2008). The 16S primer set used for other Polydesmida were not consistently successful for *Pseudopolydesmus*, and a primer set for Enolase in Diplopoda was not available, therefore custom primer sets for each gene were designed. This was accomplished by using a published transcriptome of *Pseudopolydesmus* (Rodriguez et al., 2018) and performing a local nucleotide BLAST search in the program Geneious to identify the 16S and Enolase gene regions. Primers were then developed for these regions. For 16S, a new primer set was designed (both listed in 5’ to 3’ orientation): a forward primer J-P16S (CTTTTCTCGAATAATTTTCGGGCTAA) and a reverse primer N-P16S (CGCTGTTATCCCTACAGTAGTT). Polymerase chain reaction procedures followed that of Means & Marek (2017) for 12S - 16S (5’ end), with an annealing temperature of 52 °C for 30 seconds. For Enolase, the following primer pair was designed: LR-J-PseEno (GCTGTTCCGAGTGGCGCTTC) and SR-N-PseEno (GGCACACTTGCCGTGAAAGC), with an annealing temperature of 53 °C for 30 seconds. This annealing temperature was determined after calculating an estimated melting temperature of the primers and empirically determining the optimal annealing temperature by using a gradient block to test temperature regimes between 51°C and 61°C.

Chromatograms were trimmed, bases called, and overlapping fragments assembled into contiguous sequences with the program Mesquite version 3.6 using phred and phrap in the Chromaseq version 1.51 module (Maddison and Maddison, 2010; Ewing et al., 1998). The program Prank version 170427 (Löytynoja and Goldman, 2005) was used for sequence alignment using the default gap opening and extension probabilities. Aligned gene sequences were subsequently partitioned by gene, codon position, and intron and exon locations in Mesquite. The aligned sequences used in the phylogenetic analysis have been deposited in GenBank, and a complete list of specimens and associated accession numbers are provided in Supplementary Material A. Partitions were evaluated in PartitionFinder2 to determine a best-fit partitioning model and identify appropriate models of nucleotide evolution (Lanfear et al., 2017). A concatenated matrix of the sequences was exported and used in the program RAxML version 8.2.12 (Stamatakis, 2014) to infer a phylogeny from 296 specimens using the GTRGAMMA model with the rapid bootstrap algorithm and 1,000 bootstraps. Single gene RAxML trees were inferred using the same methodology.

An integrative framework combining morphological and phylogenetic data for species delimitation in *Pseudopolydesmus* was used, according to the methods outlined in Derkarabetian & Hedin (2014, see their fig. 1). These methods involve using morphological data to delimit species clusters by using a modified PCA, and assessing these clusters with genetic data to subsequently use these clusters as *a priori* species hypotheses in a species tree, which is then validated with the program. A series of 15 morphological characters were measured for males and females of each species (Supplementary Material B) for analysis with a modified PCA described in Ezard et al. (2010): body length, head width, collum width, collum length, left 3rd antennomere length, gnathochilarium width, gnathochilarium length, clypeus width, interantennal space width, metazonite 6 width, metazonite 6 length, left leg 6 tarsus length, epiproct width, epiproct length, gonopod aperture width (males only), and cyphopod aperture width (females only). Specimens were chosen from across the distribution of each species, with preference given to specimens with all chosen characters intact. The selected specimens were chosen from collection localities that maximized geographic coverage for each species, with a maximum of 12 specimens used for each species, depending on available specimens. Analysis was done in R version 3.6.2 (Filzmoser et al., 2018; Filzmoser & Gschwandtner, 2018; Oksanen et al., 2019; R Core Team, 2019; Scrucca et al., 2016; Warnes et al., 2020), and the Ezard et al. (2010) protocol was implemented with a script written by Derkarabetian (2018). Species hypotheses informed by the RAxML-inferred phylogeny and the modified PCA results were then input into BEAST v2.6.2 (Bouckaert et al., 2019) using the *BEAST algorithm (Barido-Sottani et al., 2017, Heled & Drummond, 2010), and two species in the genus *Polydesmus* were used as outgroups to root the tree. Analyses were run for 350 million generations and logged every 10,000 generations. Resulting runs were assessed for stationarity and effective sample size (ESS) values above 200 with Tracer 1.5 (Rambaut et al., 2018). TreeAnnotator was used to reconstruct species trees from 350,000 trees (10% burnin).

## Results

A total of 1,148 *Pseudopolydesmus* specimens were collected during field work in 20 U. S. states. All known species of *Pseudopolydesmus* were collected, and these specimens were the basis of the genetic analysis. The two most commonly collected species were *P. serratus* and *P. canadensis*, which are also the most widespread species in the genus. The new specimen records were included with 2,314 unique records from museum collections and literature sources into a Darwin Core standard database of 3,462 records covering all described *Pseudopolydesmus* species (Supplementary Material C).

The phylogeny inferred by the RAxML analysis with all five genes is given in Fig. 3, and a pruned phylogeny showing only the *Pseudopolydesmus* specimens is given in Fig. 4. Each species of *Pseudopolydesmus* had bootstrap support greater than or equal to 90, and higher relationships within the genus had bootstrap support of at least 77. Within each species clade, bootstrap support among specimens varied widely: almost all specimens in the *Pseudopolydesmus paludicolus*, *P. minor,* and *P. caddo* had highly supported branches over 70, while in the *P. serratus* clade, branch support ranged from 2 to 100. *Pseudopolydesmus pinetorum*, *P. erasus, P. collinus*, and *P. canadensis* also had widely variable branch support.

**Figure 3.**
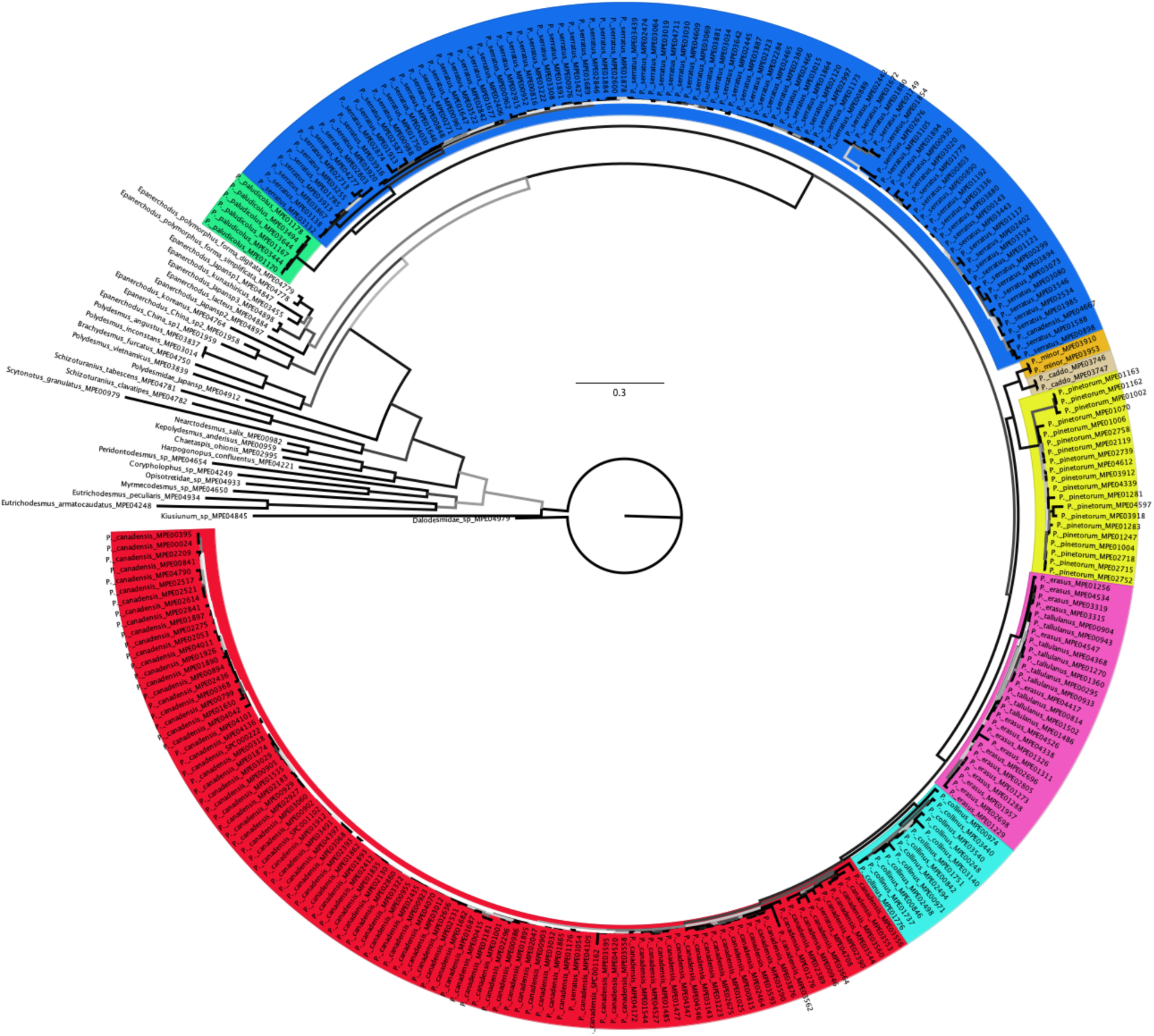
Phylogeny inferred by the RAxML analysis. Black branches indicate bootstrap support ≥ 70. Highlighted areas indicate *Pseudopolydesmus* species: green, *P. paludicolus*; blue, *P. serratus*; gold, *P. minor*; tan, *P. caddo*; yellow, *P. pinetorum*; magenta, *P. erasus*; cyan, *P. collinus*; red, *P. canadensis*. Scale: 0.3 site substitutions.

**Figure 4.**
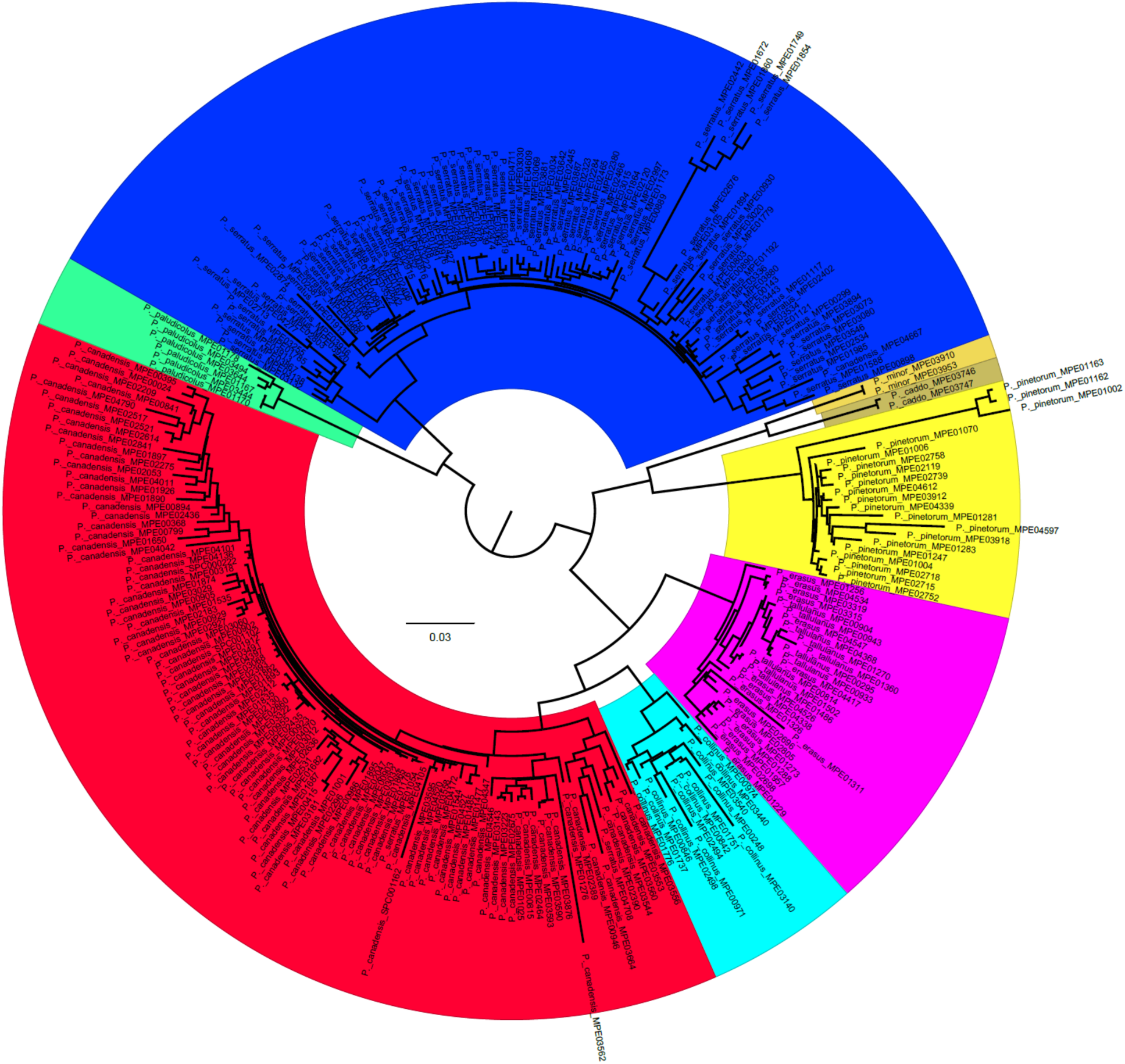
Phylogeny inferred by the RAxML analysis, only *Pseudopolydesmus* included. Highlighted areas indicate species: green, *P. paludicolus*; blue, *P. serratus*; gold, *P. minor*; tan, *P. caddo*; yellow, *P. pinetorum*; magenta, *P. erasus*; cyan, *P. collinus*; red, *P. canadensis*. Scale: 0.3 site substitutions.

The most closely-related polydesmid genus to *Pseudopolydesmus* was found to be *Epanerchodus*, a species-rich east Asian genus, but the relationship had low bootstrap support of 41. Within *Pseudopolydesmus*, the species taxonomy of Sierwald et al. (2019) was supported, with each species clade having bootstrap support of over 70%. *Pseudopolydesmus paludicolus* and *P. serratus* were found to be sister species and formed a separate clade from the rest of the species. The other species formed two smaller clades. One, composed of *P. caddo, P. minor*, and *P. pinetorum*, had *P. caddo* and *P. minor* as sister species, with *P. pinetorum* outside the *P. caddo* and *P. minor* clade. The second clade was composed of *P. erasus, P. canadensis*, and *P. collinus*. The species, *P. canadensis* and *P. collinus*, were found to be sister species, with *P. erasus* most closely related to them. Monophyly of *Pseudopolydesmus* had a high branch support of 100. The included *Epanerchodus* species had branch support values ranging between 30 and 100, and the genus was not found to be monophyletic. *Epanerchodus* was split into two clades, one comprised of mostly species found in Japan and one with species found in China and eastern Russia.

Outside of the *Pseudopolydesmus* and *Epanerchodus* clades, was a well-supported clade (bootstrap value of 98) including species of *Polydesmus* and *Brachydesmus*. *Polydesmus inconstans* Latzel, 1883 and *P. angustus* Latzel, 1884 were found to be sister taxa, and these species were sister to *Brachydesmus furcatus* Lohmander, 1936 (all bootstrap values of 100).

Sister to this clade was *Polydesmus vietnamicus* Anh, 2009 (bootstrap value of 68). Sister to this clade and the *Pseudopolydesmus* and *Epanerchodus* clade was an unidentified species of Polydesmidae from Japan (bootstrap value of 87). Sister to the rest of the Polydesmidae was a clade with *Scytonotus granulatus* (Say, 1821) sister to *Schizoturanius clavatipes* (Stuxberg, 1876) and *Schizoturanius tabescens* (Stuxberg, 1876) (all bootstrap values greater than or equal to 77).

The other Polydesmidea families included in the analysis had generally high support values within each family, but higher relationships among the families were not well supported. Macrosternodesmidae was the sister group to the Polydesmidae, but with a bootstrap value of 42, while bootstrap values between species in the family were 99 and 100. Macrosternodesmidae and Polydesmidae were sister to the rest of the Polydesmidea families, with bootstrap support of 43. Opisotretidae was sister to an unidentified species of *Peridontodesmus* Silvestri, 1896 in the Cryptodesmidae (bootstrap support of 75). This clade was sister to a clade of the Haplodesmidae and Pyrgodesmidae (bootstrap support of 40 and bootstrap support of 53 for Haplodesmidae + Pyrgodesmidae). An unidentified species of *Kiusiunum* Verhoeff, 1942 was found to be sister to the rest of these clades, therefore this analysis did not find the Cryptodesmidae to be monophyletic.

For the modified PCA, a total of 69 male *Pseudopolydesmus* specimens were measured for 15 characters. Six morphological clusters were identified, and two principal components were retained (Fig. 5). Visualization of the eigenvalues with a scree plot showed that the first principal component (body length) explained about 96% of the variance, while the second principal component explained only about 2% of the variance. The second component (and the other measured characters) is highly positively correlated with body length. The identified morphological clusters differed from the species clades identified by the genetic analysis, and species were subsequently separated using the genetic clades to formulate the *a priori* species hypotheses for the species tree analysis.

**Figure 5.**
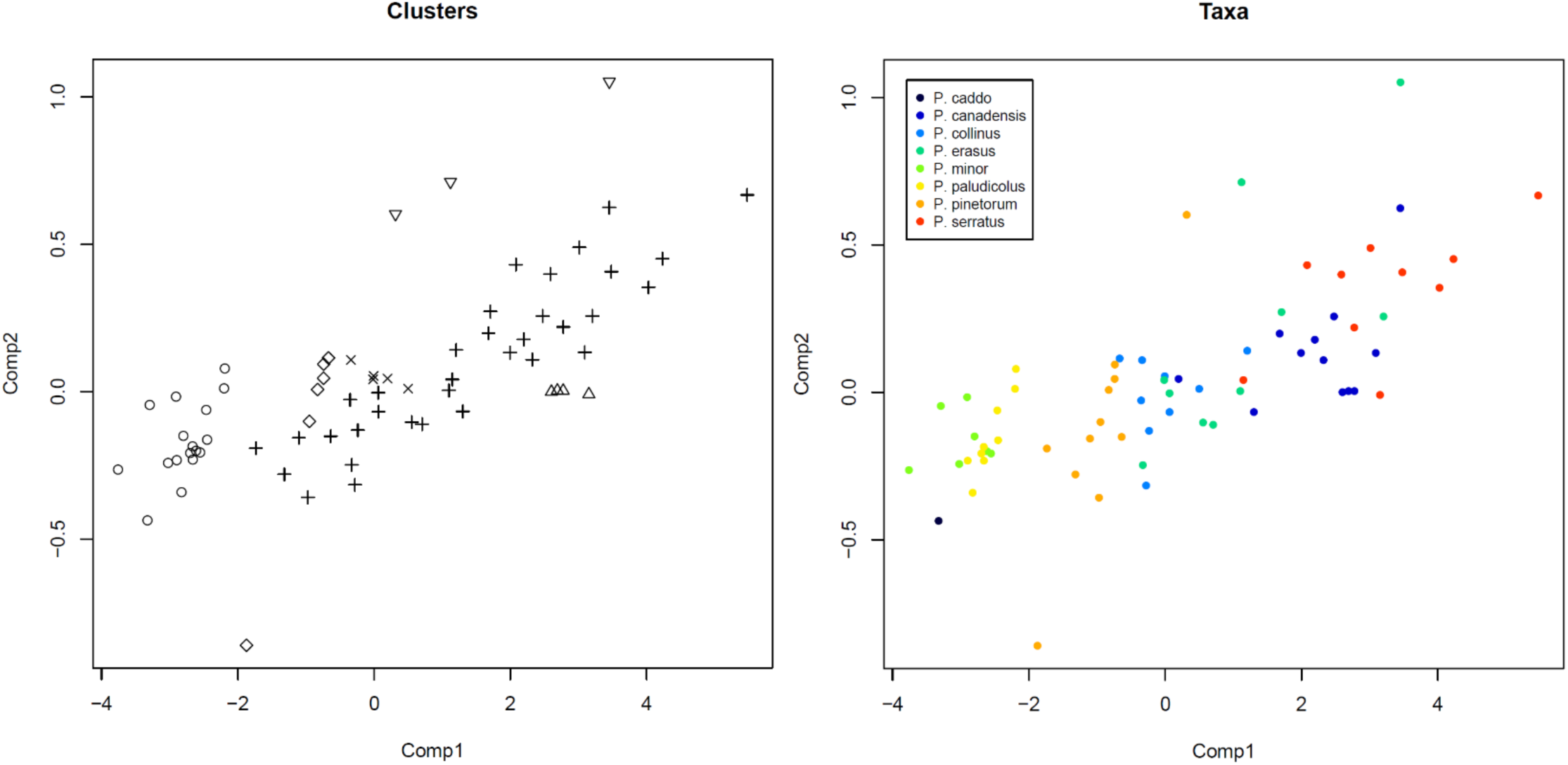
Multivariate clustering with PCA. Left panel: clusters identified by modified PCA. Right panel: *Pseudopolydesmus* specimens colored by species.

The species tree inferred by the *BEAST analysis is show in Figure 6. The relationships in the *BEAST phylogeny differ from those in the RAxML-inferred phylogeny. In the *BEAST tree, *P. caddo* and *P. minor* are indicated as sister species with high support, and form a clade with *P. pinetorum*, but this clade is sister to all the other *Pseudopolydesmus* species. In this other clade, *P. paludicolus* is sister to the other *Pseudopolydesmus* species, and *P. erasus* is sister to the other three species. The species *P. collinus* is sister to a clade of *P. canadensis* and *P. serratus*, a relationship not found in the RAxML phylogeny. This may be a result of low support values in the *BEAST phylogeny, as ESS values for this phylogeny were very low, at 13. Within the phylogeny, only the relationship between *P. caddo* and *P. minor*, and *P. erasus* being sister to the (*P. collinus*, (*P. canadensis, P. serratus*)) clade are well-supported.

**Figure 6.**
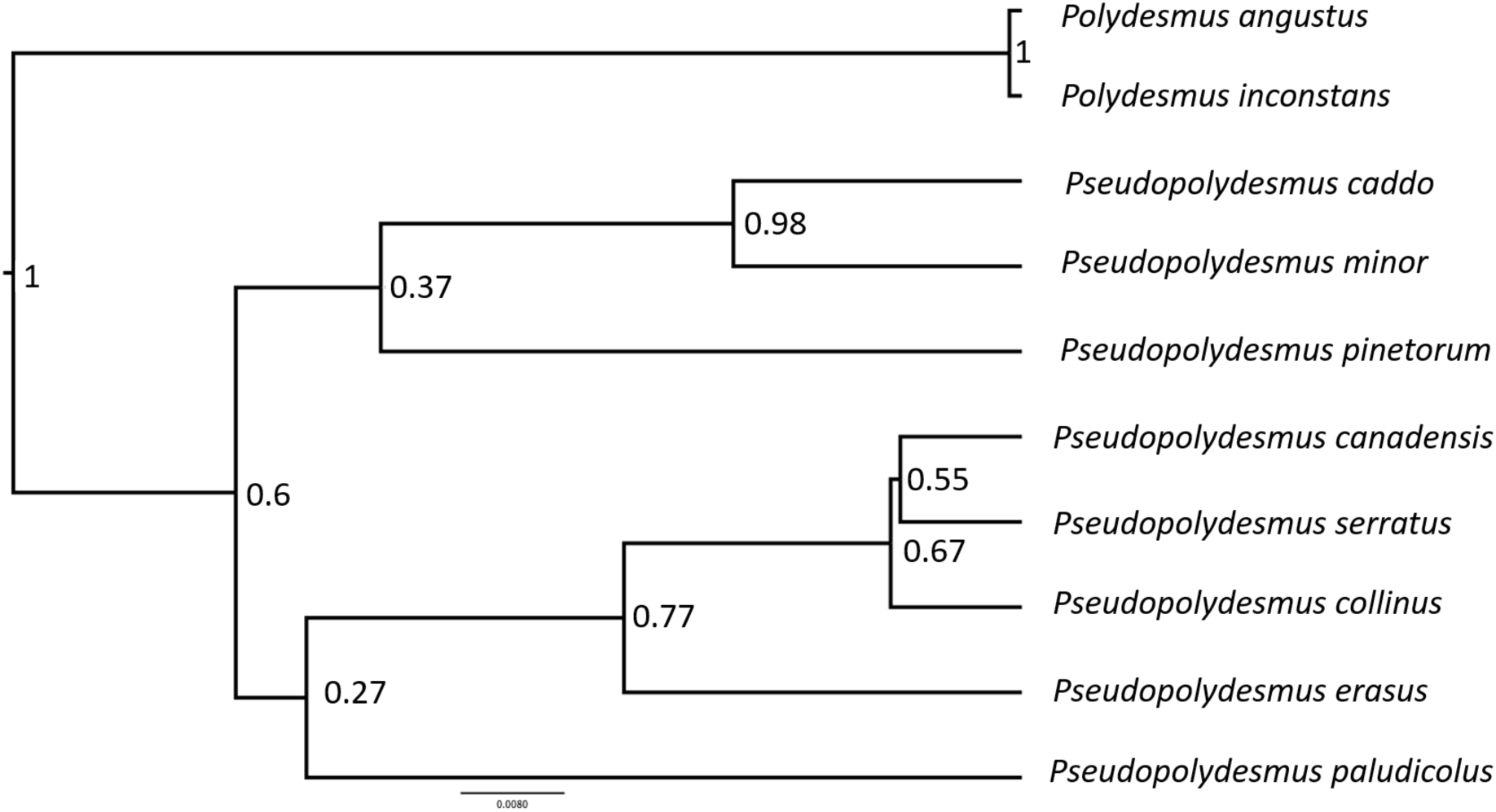
Species tree for *Pseudopolydesmus* inferred by *BEAST, with *Polydesmus* outgroups. Numbers at nodes indicate posterior probability support.

Due to the incongruence between the RAxML phylogeny and the *BEAST phylogeny, the relationships within the genus indicated by the RAxML phylogeny are followed in the Taxonomic Part of this study. This is due to the higher support values in the RAxML analysis, as well as its congruence with morphological characters.

### Taxonomic Part

Class **Diplopoda** de Blainville in Gervais, 1844

Order **Polydesmida** Pocock, 1887

Suborder **Polydesmidea** Leach, 1815

Family **Polydesmidae** Leach, 1815

Genus *Pseudopolydesmus* Attems, 1898

**Type species**: *Polydesmus canadensis* Newport, 1844.

Included species:

*Pseudopolydesmus caddo* Chamberlin, 1949

*Pseudopolydesmus canadensis* (Newport, 1844)

*Pseudopolydesmus collinus* Hoffman, 1974

*Pseudopolydesmus erasus* (Loomis, 1943)

*Pseudopolydesmus minor* (Bollman, 1888)

*Pseudopolydesmus paludicolus* Hoffman, 1950

*Pseudopolydesmus pinetorum* (Bollman, 1888)

*Pseudopolydesmus serratus* (Say, 1821)

Synonymies, morphological descriptions, gonopod images, and a key to the species of the genus can be found in Sierwald et al. (2019).

**Pseudopolydesmus caddo Chamberlin, 1949**

*Pseudopolydesmus bidens* Loomis, 1959

Vernacular name: “The Caddo Marsh Flatback”

**Figs. 7, 8.**

**Diagnosis**: Pulvillus pointed (Fig. 7). Lacking process *e1*, not present as in *P. erasus*; lacking process *e2*, not present as in all other *Pseudopolydesmus* species; process *e3* present and on a lamina, not absent as in *P. minor*; process *e4* absent, not present as in all other *Pseudopolydesmus* species; process *m1* fin-shaped, not triangular as in *P. minor*; process *m2* small, triangular, not large as in *P. minor*; process *m3* absent, not present as in *P. paludicolus*; process *m4* present, not absent as in *P. paludicolus*. Processes *m2* and *e3* connected by transverse ridge.

**Figure 7.**
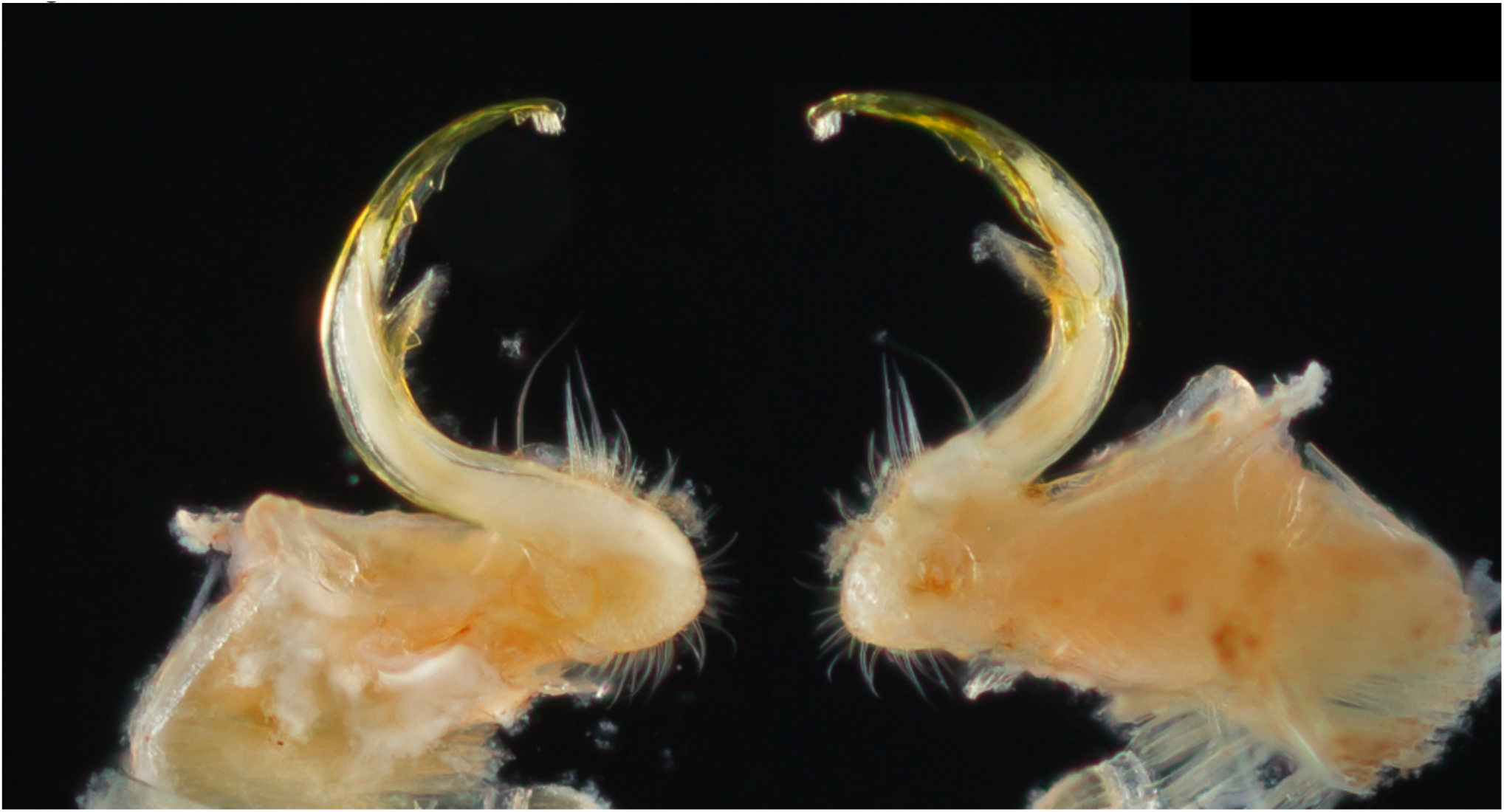
Left gonopod of *Pseudopolydesmus caddo* (MPE03746, St. Landry Parish, Louisiana), medial view (left) and ectal view (right).

**Color**: Color in life unknown, but preserved specimens dark maroon.

**Distribution**: *Pseudopolydesmus caddo* is only known from Louisiana and eastern Texas (Fig. 8).

**Figure 8.**
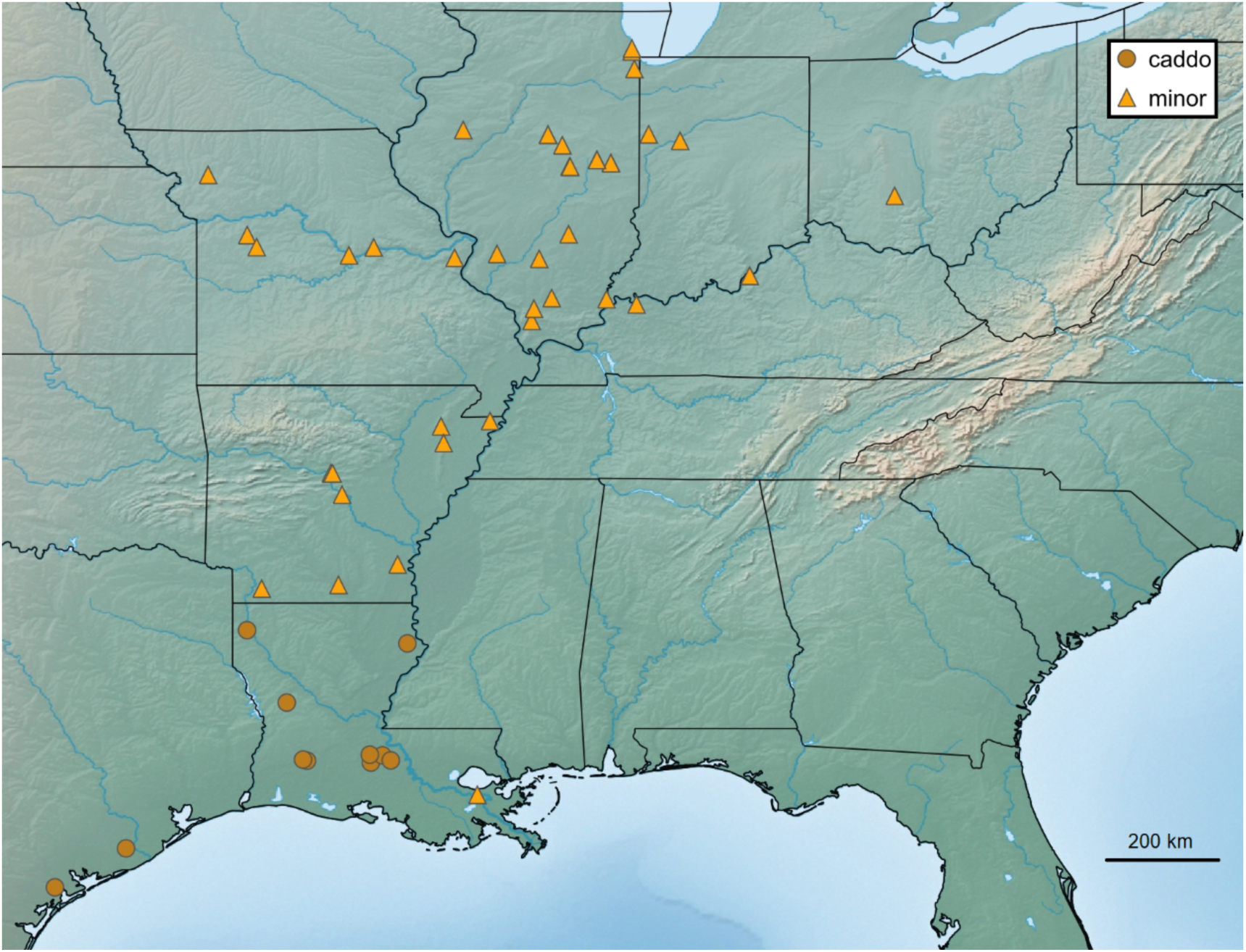
Distribution of *Pseudopolydesmus caddo* (dots) and *Pseudopolydesmus minor* (triangles) in the eastern United States.

**Ecology**: *Pseudopolydesmus caddo* is one of the three species in the genus known from swamps and floodplains (the others being *P. minor* and *P. paludicolus*). Locality and habitat data indicate it is found in areas adjacent to streams, swamps, and rivers, and it has been collected on floating wooden debris in these habitats. This species is the most poorly-known in the genus, having only been collected from about a dozen localities. It has most often been collected during Spring and late Fall.

**Pseudopolydesmus canadensis (Newport, 1844)**

*Polydesmus glaucescens* C. L. Koch, 1847

*Polydesmus nitidus* Bollman, 1887

*Polydesmus branneri* Bollman, 1887

*Polydesmus echinogon* Chamberlin, 1942

*Polydesmus conlatus* Chamberlin, 1943

*Dixidesmus sylvicolens* Chamberlin, 1943

*Dixidesmus christianus* Chamberlin, 1946

*Dixidesmus catskillus* Chamberlin, 1947

*Dixidesmus phanus* Chamberlin, 1951

*Dixidesmus gausodicrorhachus* Johnson, 1954

Vernacular name: “The Canadian Flatback”

**Figs. 2, 9-11.**

**Diagnosis:** Pulvillus pointed (Fig. 9). Process *e1* present, abruptly bent basally, larger than in *P. collinus*; processes *e2* and *e3* present and connected by a basal stalk, *e2* not a single process as in *P. serratus*; process *e4* small, not absent as in *P. caddo*; processes *m1* and *m2* present, as in all *Pseudopolydesmus* species, quadrate and triangular, respectively; process *m3* present, triangular and positioned near midline of acropodite, not absent as in *P. serratus*; process *m4* triangular, not absent as in *P. serratus*.

**Figure 9.**
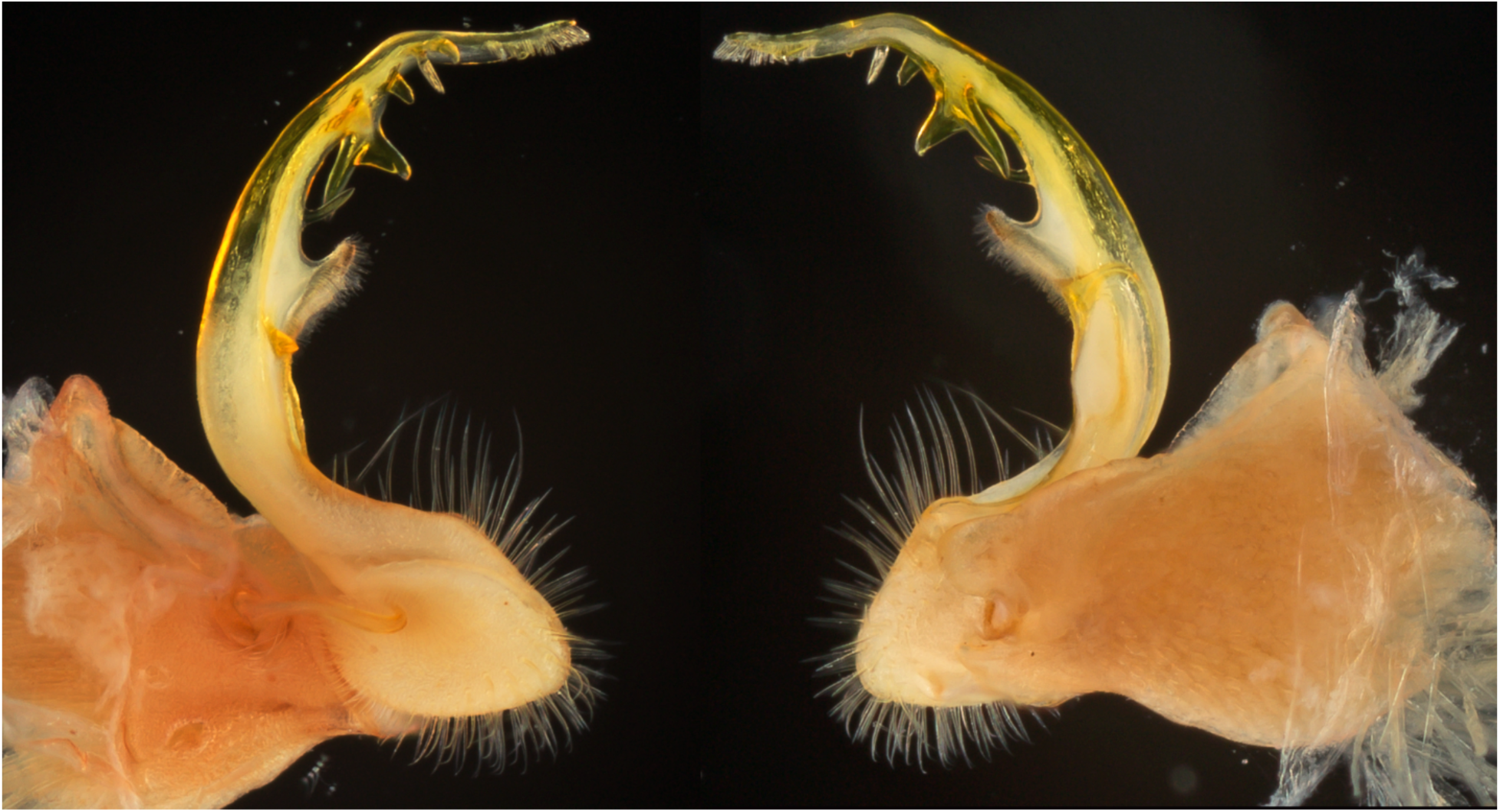
Left gonopod of *Pseudopolydesmus canadensis* (MPE00799, Craig County, Virginia), medial view (left) and ectal view (right).

**Color**: Specimens of *P. canadensis* most often have a broad longitudinal black dorsal stripe with brick-red paranota (Fig. 10), but the paranota color can sometimes be a lighter pink. Low-elevation populations in the Atlantic and Gulf Coastal Plains lack the black stripe and red paranota, and are instead light pink.

**Figure 10.**
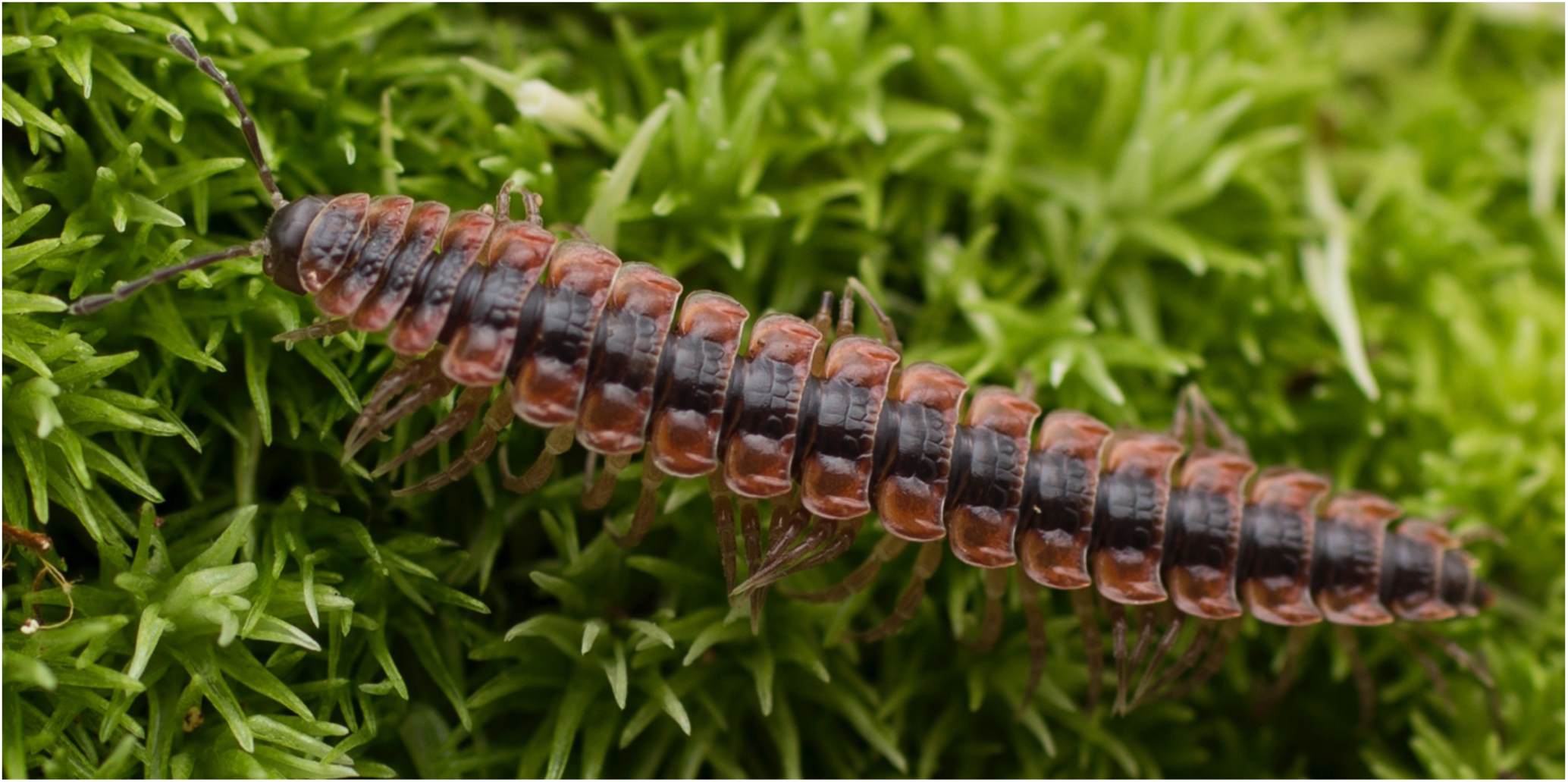
*Pseudopolydesmus canadensis,* adult male from Ashe County, North Carolina (MPE04390).

**Distribution**: *Pseudopolydesmus canadensis* is one of the most widespread species in the genus, ranging from Québec and Ontario (Canada) south to Florida (Fig. 11). It is the most abundant *Pseudopolydesmus* species in Appalachia and the northern U.S. and Canada, but also ranges southward to the Atlantic and Gulf Coastal Plains. In this southern portion of its range, body size is noticeably smaller and it lacks the bold coloration seen in the rest of its distribution. Individuals in the Coastal Plain areas have light pink tergites and no contrastingly-colored paranota.

**Figure 11.**
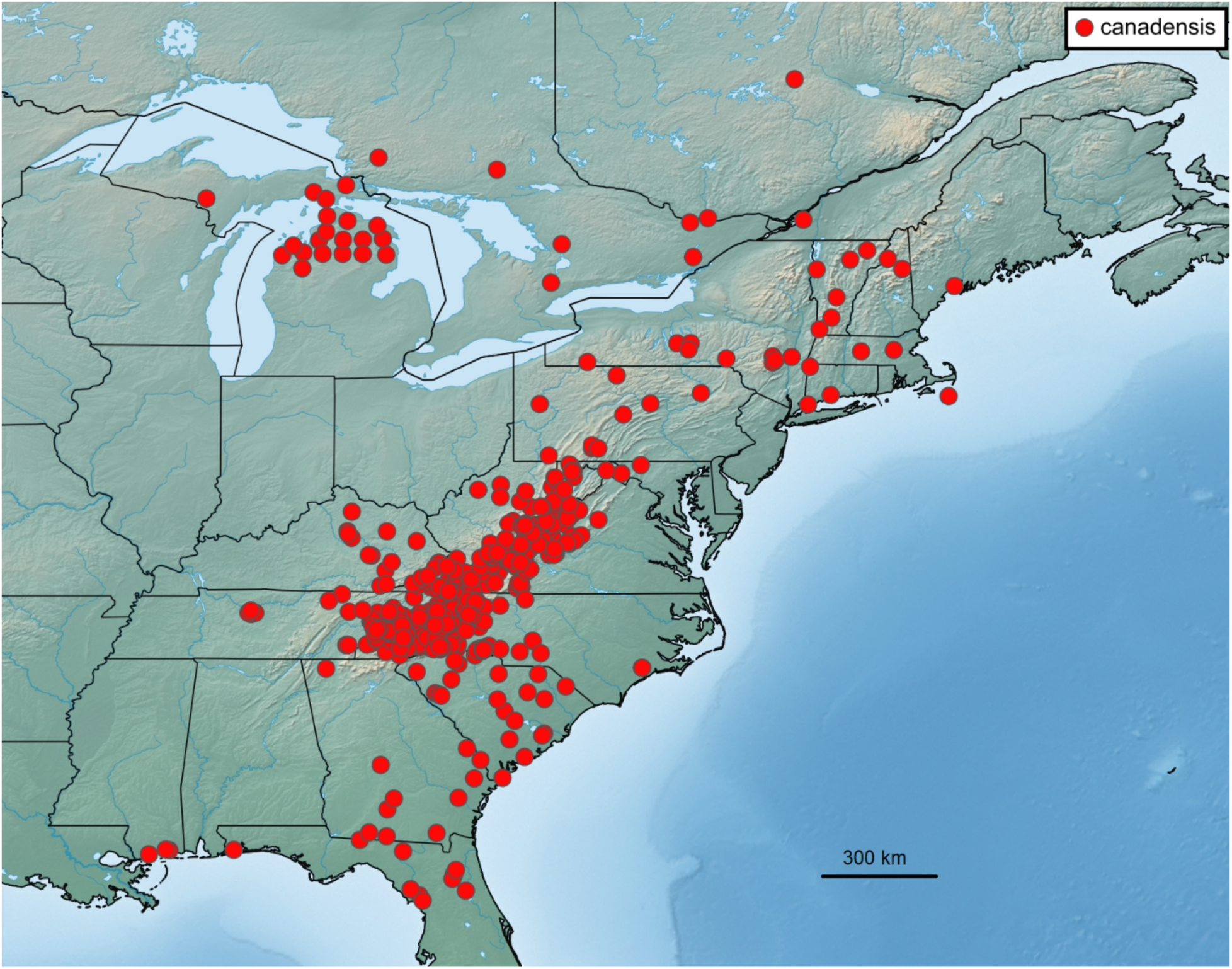
Distribution of *Pseudopolydesmus canadensis*.

**Ecology**: This species has been found in a wide range of habitats, from generalized deciduous forests, to hemlock (*Tsuga* spp.) and rhododendron (*Rhododendron* spp.) cove forests, and sandy pine (*Pinus* spp.) forests. Label data indicate it has been collected in deciduous forests of oak (*Quercus* spp.), hickory (*Carya* spp.), maple (*Acer* spp.), and beech (*Fagus grandifolia*). It has been collected at near sea level in the Coastal Plain regions, all the way up to Mount Mitchell, North Carolina, the highest peak in the Appalachians (2,037 meters).

**Pseudopolydesmus collinus Hoffman, 1974**

Vernacular name: “The Rolling Hills Flatback” **Figs. 12-14**.

**Figure 12.**
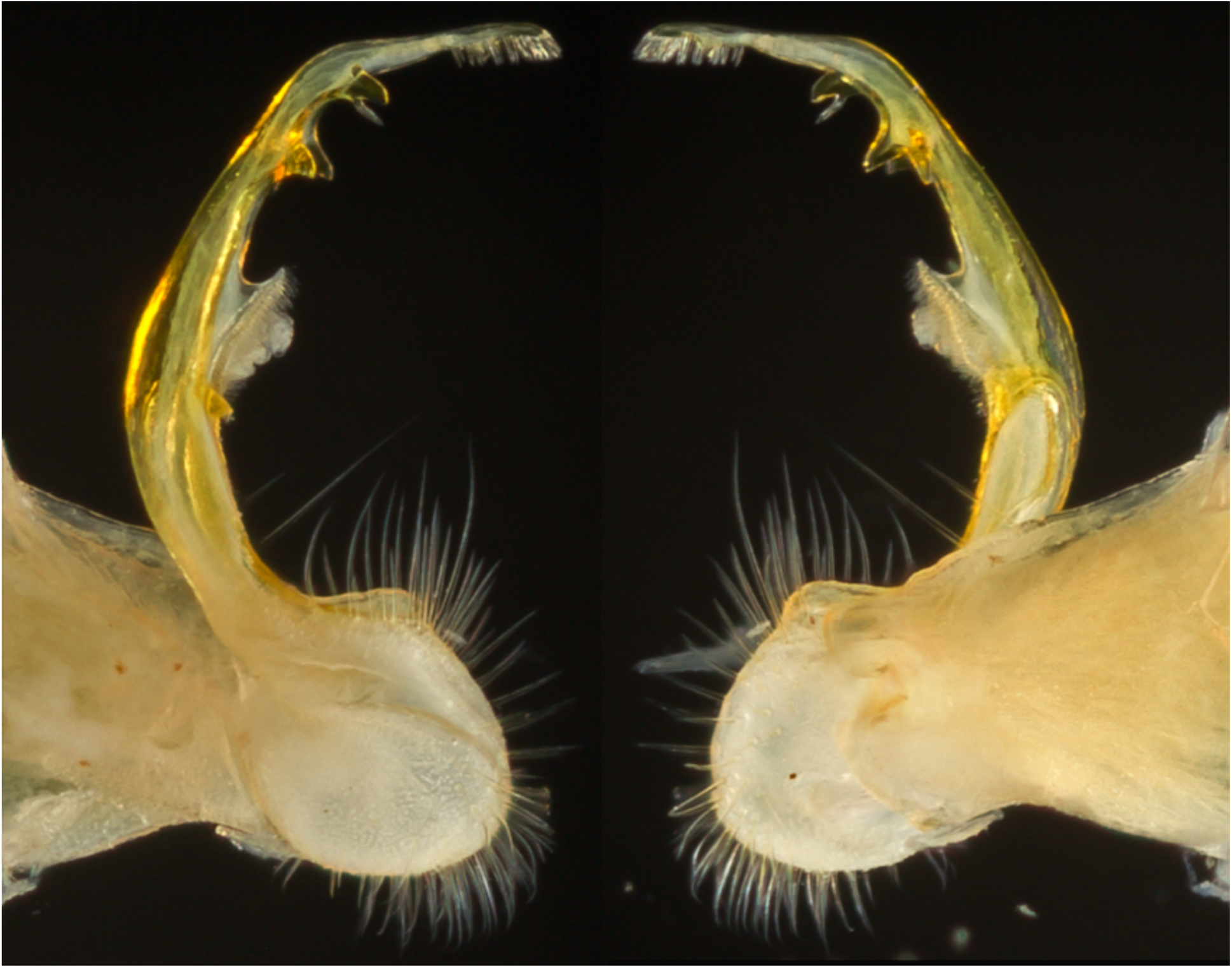
Left gonopod of *Pseudopolydesmus collinus* (MPE00842, Patrick County, Virginia), medial view (left) and ectal view (right).

**Figure 13.**
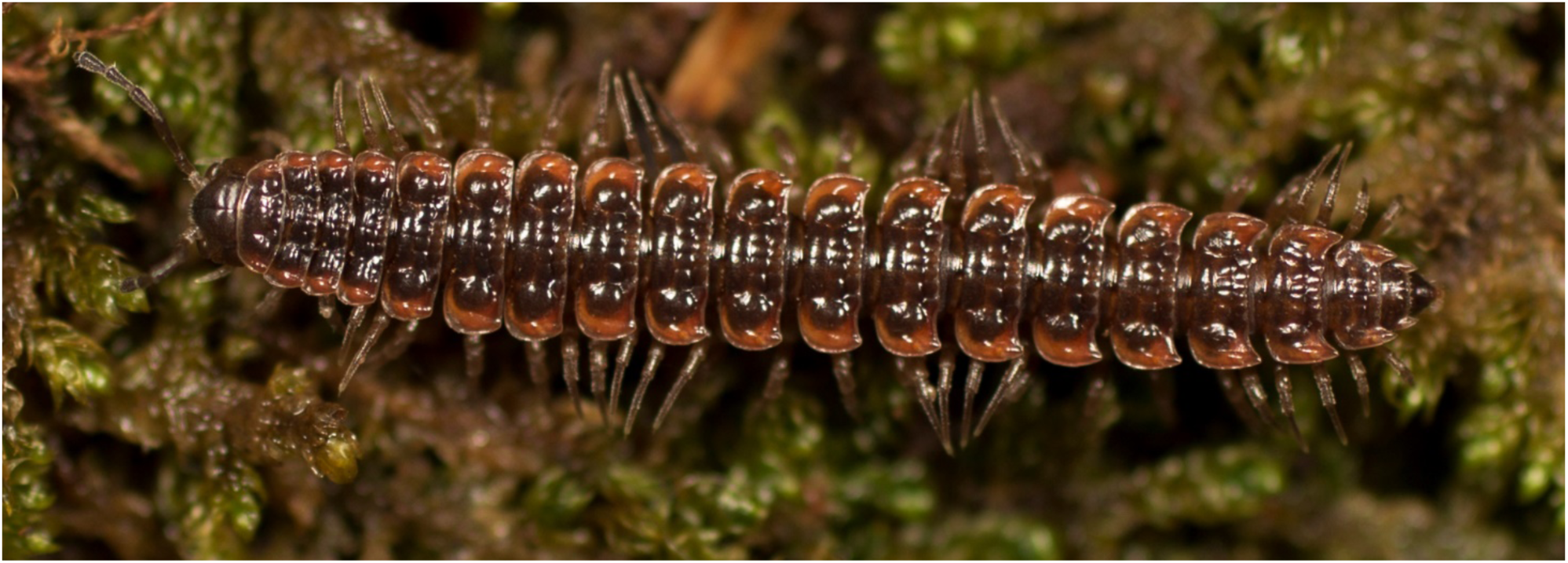
*Pseudopolydesmus collinus*, dorsal view of adult male from Carroll County, Virginia (VTEC, MPE03520).

**Figure 14.**
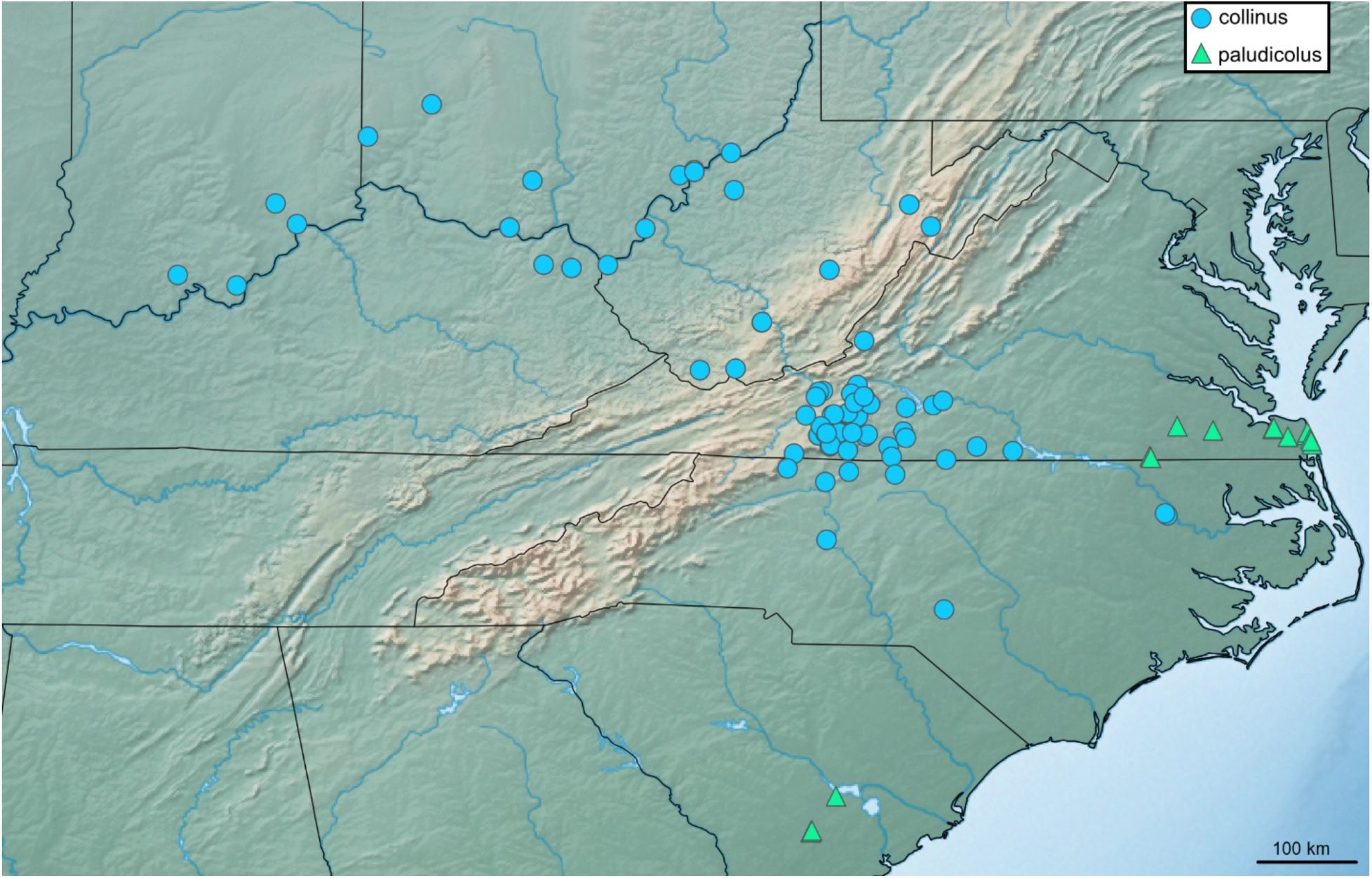
Distribution of *Pseudopolydesmus collinus* (dots) and *P. paludicolus* (triangles).

**Diagnosis**: Pulvillus pointed, slightly recurved (Fig. 12). Process *e1* small, straight, pointed, sometimes absent, not large as in *P. canadensis*; processes *e2* and *e3* connected by basal stalk, smaller than in *P. canadensis*, process *e2* not absent as in *P. serratus*; process *e4* triangular with a broad base, not absent as in *P. caddo*; process *m1* spade-shaped, not broadly quadrate as in *P. canadensis*; *m2* fin-shaped, not triangular as in *P. canadensis*; process *m3* absent, not present as in *P. canadensis*; process *m4* large, triangular, not absent as in *P. serratus*.

**Color**: *Pseudopolydesmus collinus* has light pink to red paranota edges, with the rest of the tergites being a dark maroon to light pink color (Fig. 13).

**Distribution**: *Pseudopolydesmus collinus* is known from six states: the southern portions of Indiana and Ohio and adjacent northern Kentucky, south through West Virginia, Virginia, and North Carolina (Fig. 14).

**Ecology**: This species is often found near rivers and streams at low to middle elevations, being recorded from elevations spanning from 169 m to 1109 m. Label data indicates it is most often found in deciduous forests of oak (*Quercus* spp.), maple (*Acer* spp.), tulip tree (*Liriodendron tulipifera)*, American beech (*Fagus grandifolia*), witch hazel (*Hamamelis virginiana*), and buckeye (*Aesculus* spp.). It has also been collected in hemlock (*Tsuga* spp.) and rhododendron (*Rhododendron* spp.) coves.

**Pseudopolydesmus erasus (Loomis, 1943)**

*Dixidesmus tallulanus* Chamberlin, 1943

*Dixidesmus penicillus* Chamberlin, 1943

*Dixidesmus humilidens* Chamberlin, 1943

Vernacular name: “The Southern Flatback” **Figures 15-17**.

**Figure 15.**
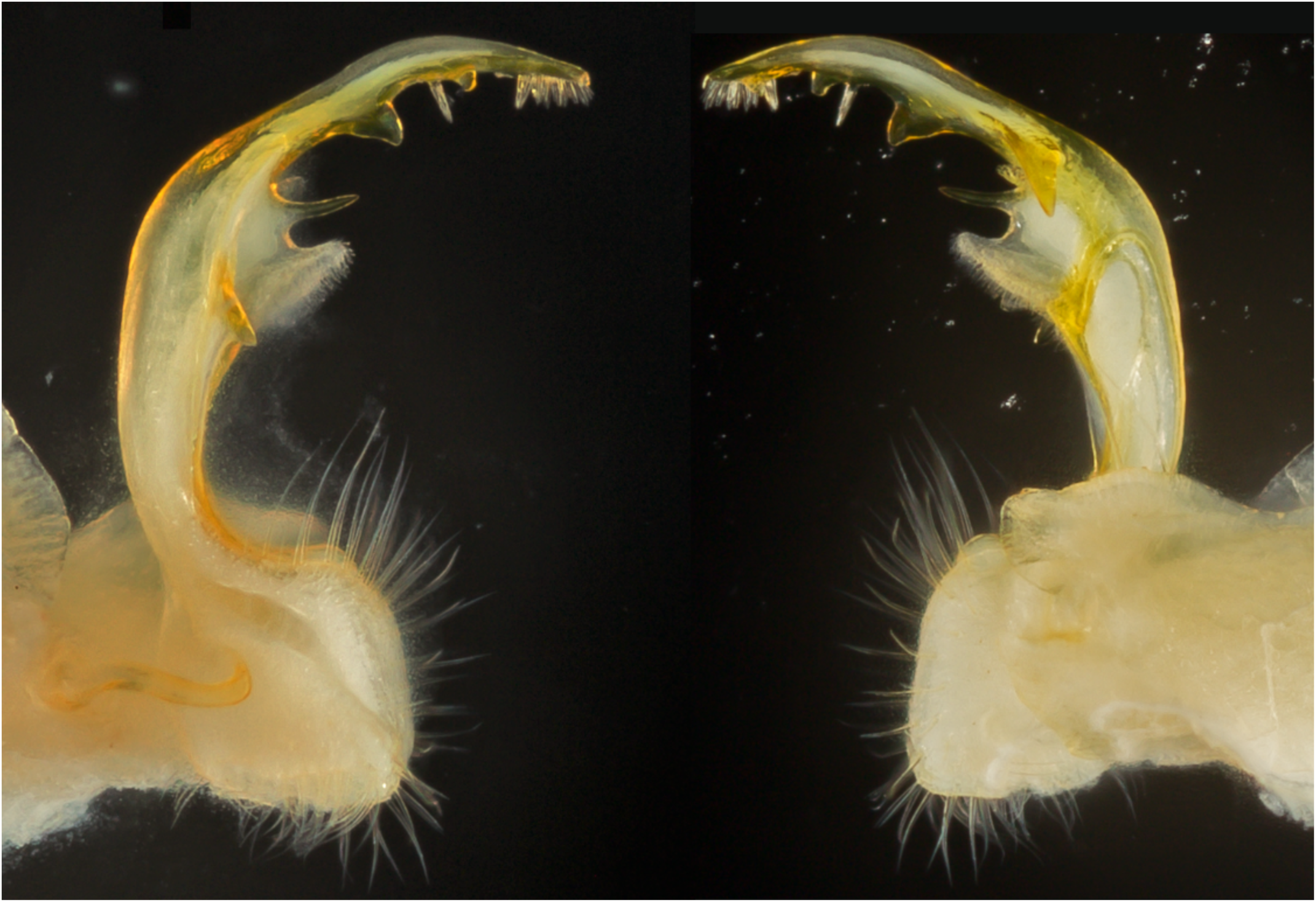
Left gonopod of *Pseudopolydesmus erasus* (MPE01326, Putnam County, Tennessee), medial view (left) and ectal view (right).

**Figure 16.**
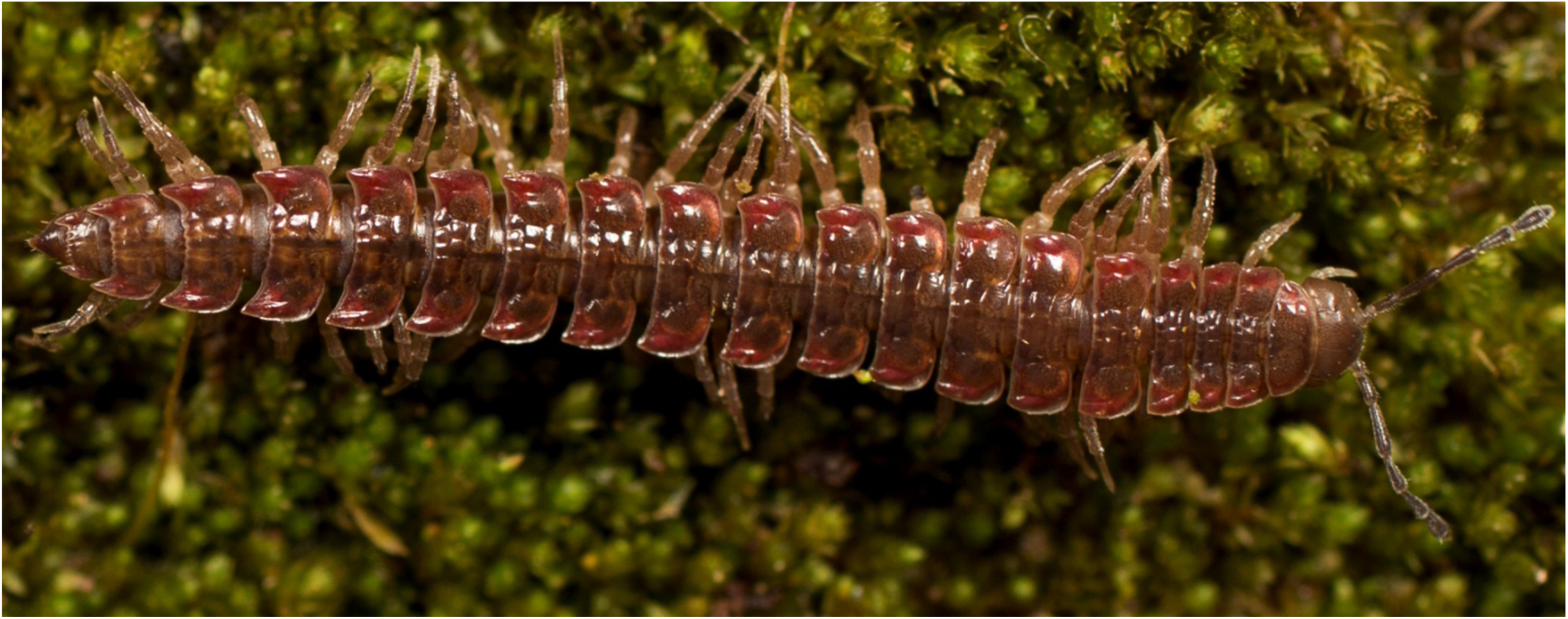
*Pseudopolydesmus erasus*, dorsal view of male from Van Buren County, Tennessee (MPE01229).

**Figure 17.**
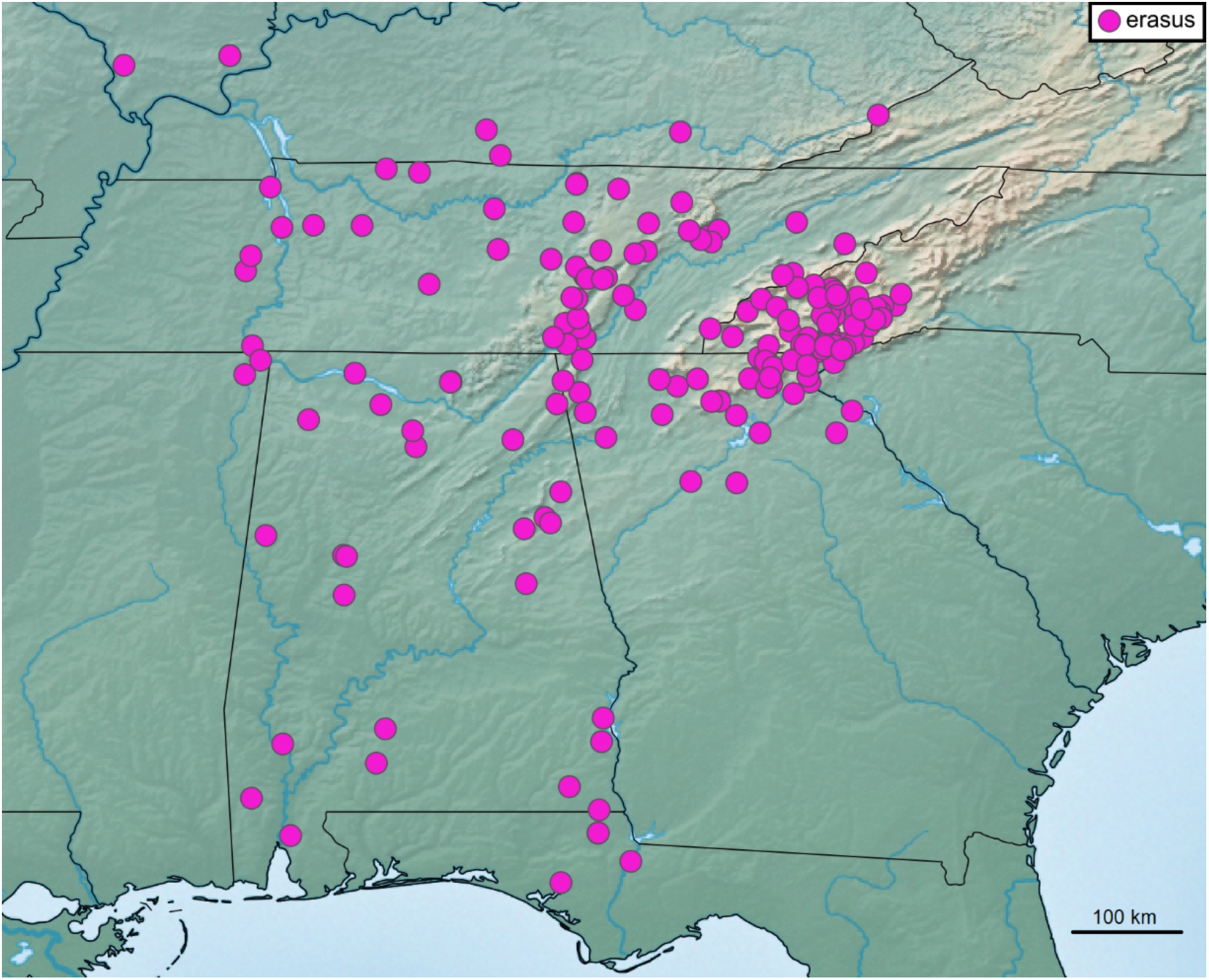
Distribution of *Pseudopolydesmus erasus*.

**Diagnosis**: Pulvillus large, rounded distally (Fig. 15). Process *e1* pointed and straight, arising from transversely expanded area, not bent at base as in *P. canadensis*; process *e2* large, triangular, not connected to process *e3* by basal stalk as in *P. canadensis*; process *e3* small to medium sized and triangular, sometimes absent; process *e4* small, triangular, not absent as in *P. caddo*; process *m1* large, pointed to spade-shaped; process *m2* triangular, not fin-shaped as in *P. collinus*; process *m3* absent, not present as in *P. canadensis*; process *m4* triangular, with rounded apex, not absent as in *P. serratus*.

**Color**: *Pseudopolydesmus erasus* shows some color variability, but tends to have tergites mostly maroon-brown with paranota edges pink to light red, or sometimes light orange (Fig. 16).

**Distribution**: *Pseudopolydesmus erasus* is found throughout the southern United States, ranging from the southern tip of Illinois south to the Florida panhandle (Fig. 17). It is the most common *Pseudopolydesmus* species throughout much of its range, but is sympatric with *P. canadensis, P. minor, P. pinetorum*, and *P. serratus* in portions of its range.

**Ecology**: *Pseudopolydesmus erasus* is a general forest species, and can be found in mixed deciduous woods, pine woods, and hemlock-rhododendron coves. It occurs in a variety of habitats throughout its range, from low elevation plains to high elevation mountain forests. It is not particularly tied to streams or swamps, and can tolerate drier habitats.

**Pseudopolydesmus minor (Bollman, 1888)**

Polydesmus neoterus Chamberlin, 1942

*Polydesmus euthetus* Chamberlin, 1942

Vernacular name: “The Small Swamp Flatback”

**Figs. 8, 18-19.**

**Diagnosis**: Pulvillus pointed and elongate (Fig. 18). Process *e1* absent, not present as in *P. erasus*; process *e2* small, triangular, on ectal flange, not absent as in *P. caddo*; process *e3* absent, with the occurrence of an ectal flange in its place, not triangular as in *P. caddo*; process *e4* small, sharp, not absent as in *P. caddo*; process *m1* large, triangular, not fin-shaped as in *P. caddo*; process *m2* large, triangular, not small as in *P. caddo*; process *m3* absent, not present as in *P. paludicolus*; process *m4* triangular, not absent as in *P. paludicolus*.

**Figure 18.**
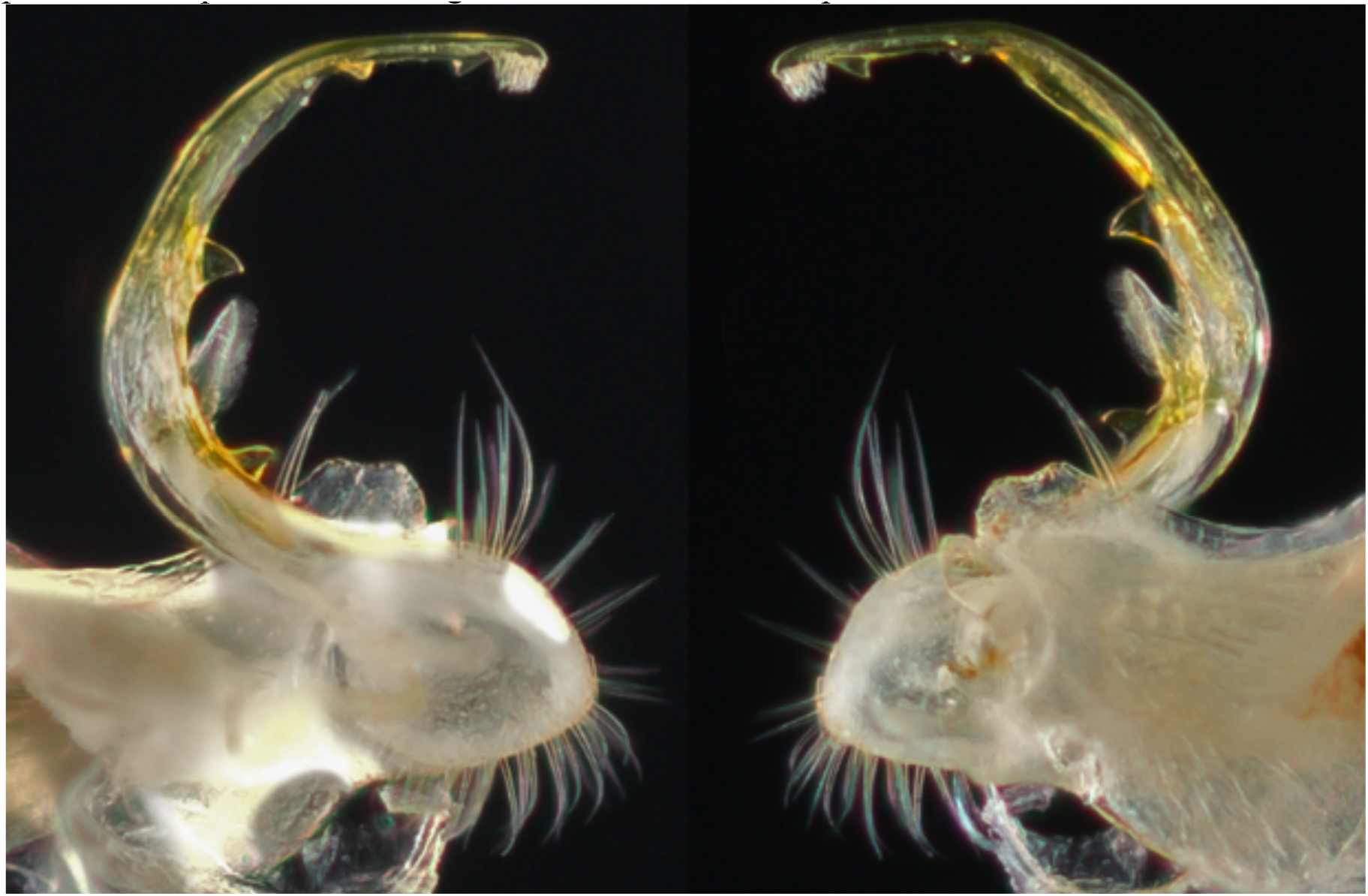
Left gonopod of *Pseudopolydesmus minor* (MPE03910, Champaign County, Illinois), medial view (left) and ectal view (right).

**Color**: *Pseudopolydesmus minor* is a small species with dark brown tergites and paranota edges light tan-brown (Fig. 19).

**Figure 19.**
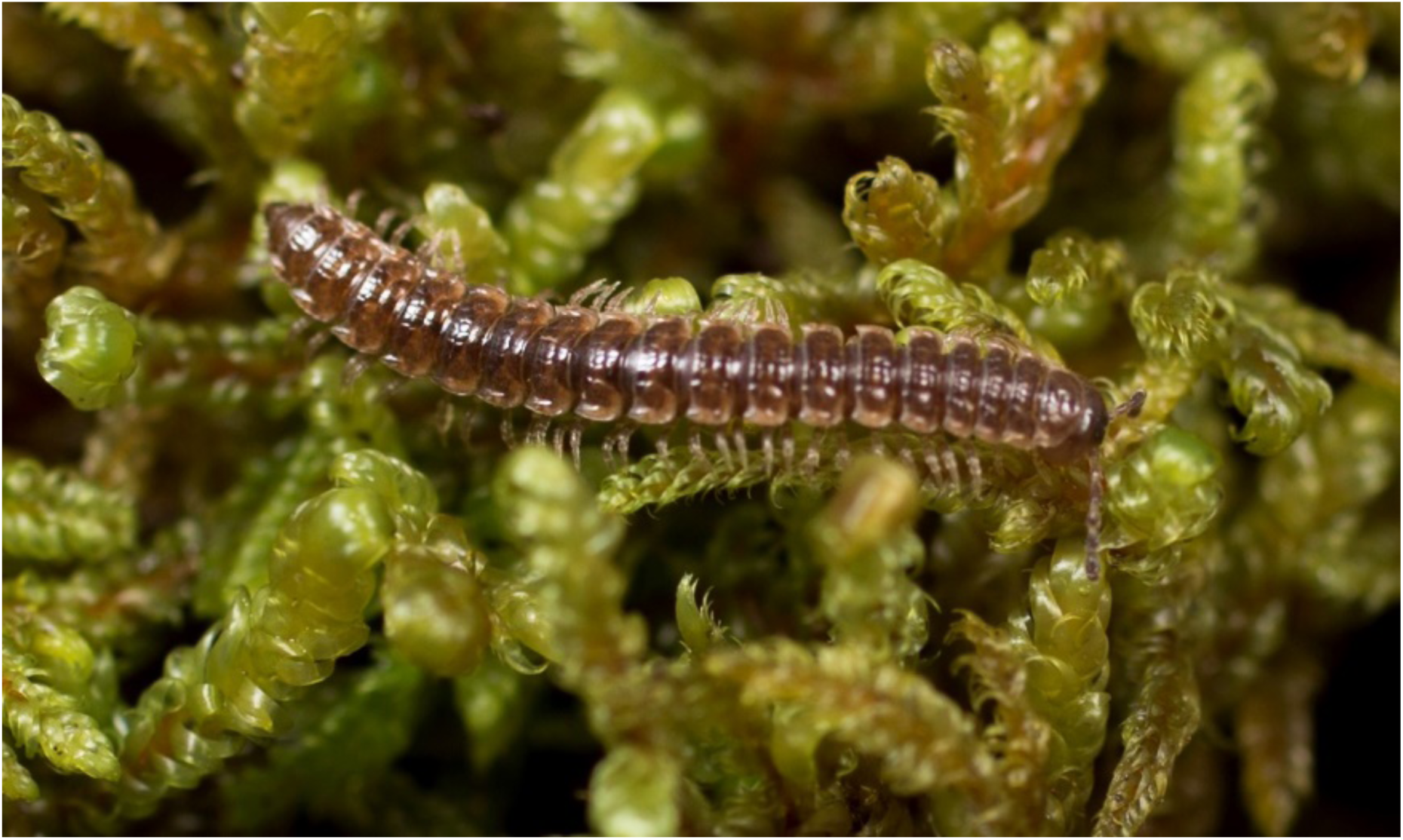
*Pseudopolydesmus minor*, dorsal view of adult female from Piatt County, Illinois (MPE03923).

**Distribution**: This species is the most widespread of all the wetland-adapted *Pseudopolydesmus* species (the others being *P. caddo* and *P. paludicolus*), and is found from Louisiana north to Indiana, mainly along the Mississippi River drainage (Fig. 8). Two records from far outside the core of the distribution are included, one from New Orleans, Louisiana, and another from Washington Court House, Ohio.

**Ecology**: *Pseudopolydesmus minor* is a swamp-adapted species found in floodplains and swamps associated with rivers and streams. Label data indicates it is associated with deciduous floodplain forests composed of oaks (*Quercus* spp.), maples (*Acer* spp., including *A. negundo* and *A. saccharinum*), hickory (*Carya* spp.), and American sycamore (*Platanus occidentalis*).

Within these habitats, *P. minor* is often found associated with stumps in grassy areas, or under logs and other woody debris, particularly after recent flooding events.

***Pseudopolydesmus paludicolus Hoffman*, 1950**

Vernacular name: “The Virginia Swamp Flatback” **Figs. 14, 20-21.**

**Diagnosis**: Pulvillus small, pointed (Fig. 20). Process *e1* absent, not present as in *P. canadensis*; process *e2* small, pointed, not large as in *P. serratus*; process *e3* absent, not present as in *P. caddo*; process *e4* large, triangular, slightly curved distally, not small as in *P. serratus*; process *m1* small, spade-shaped, not large as in *P. erasus*; process *m2* triangular, connected to process *m3* by lamina, both processes medium-sized, process *m2* not connected to *e2* by a transverse ridge as in *P. serratus*, process *m3* not absent as in *P serratus*; process *m4* absent, not present as in *P. caddo*.

**Figure 20.**
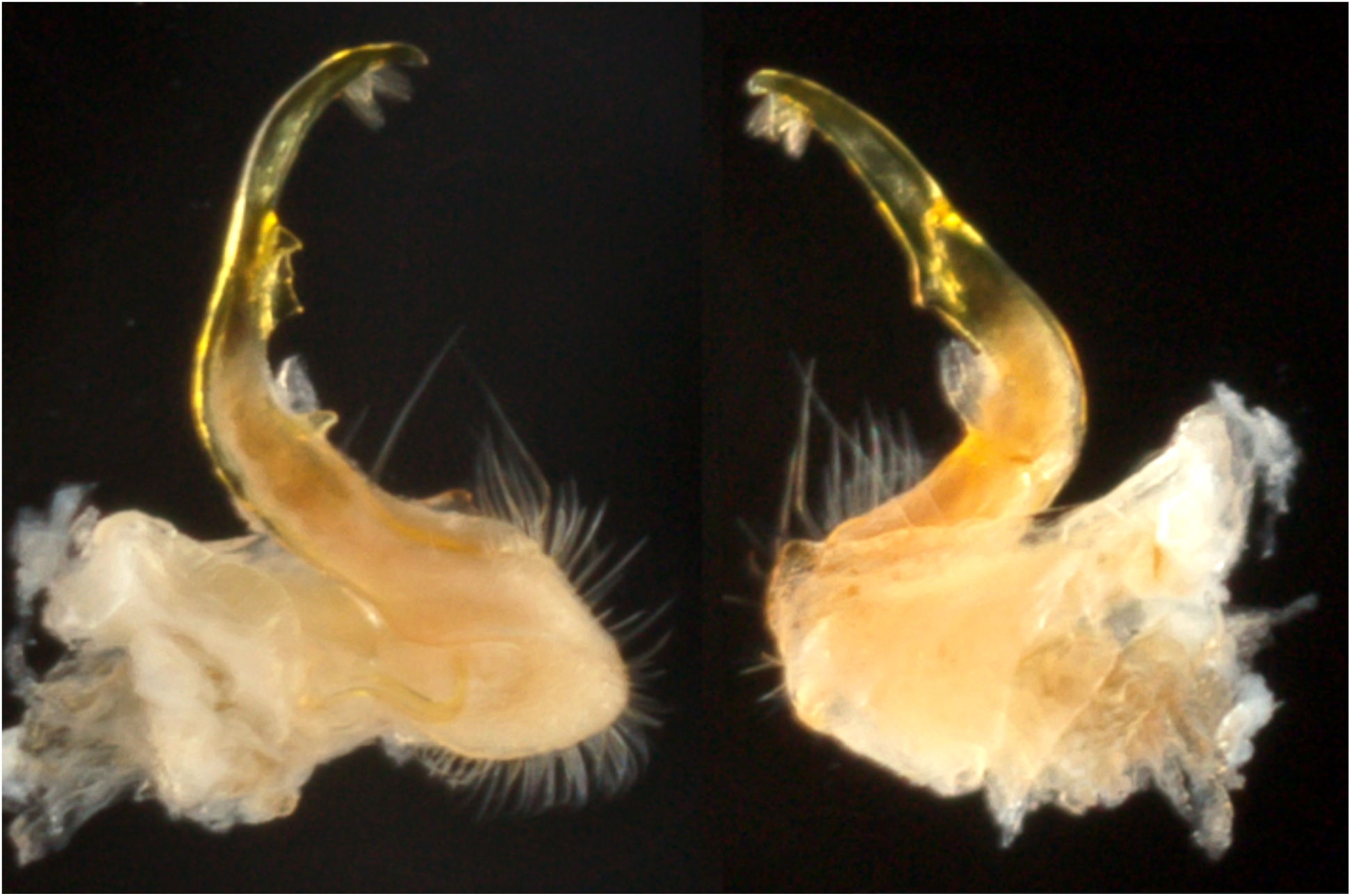
Left gonopod of *Pseudopolydesmus paludicolus* (MPE01169, Greensville County, Virginia), medial view (left) and ectal view (right).

**Color**: Maroon-brown tergites with paranota the same color (Fig. 21).

**Figure 21.**
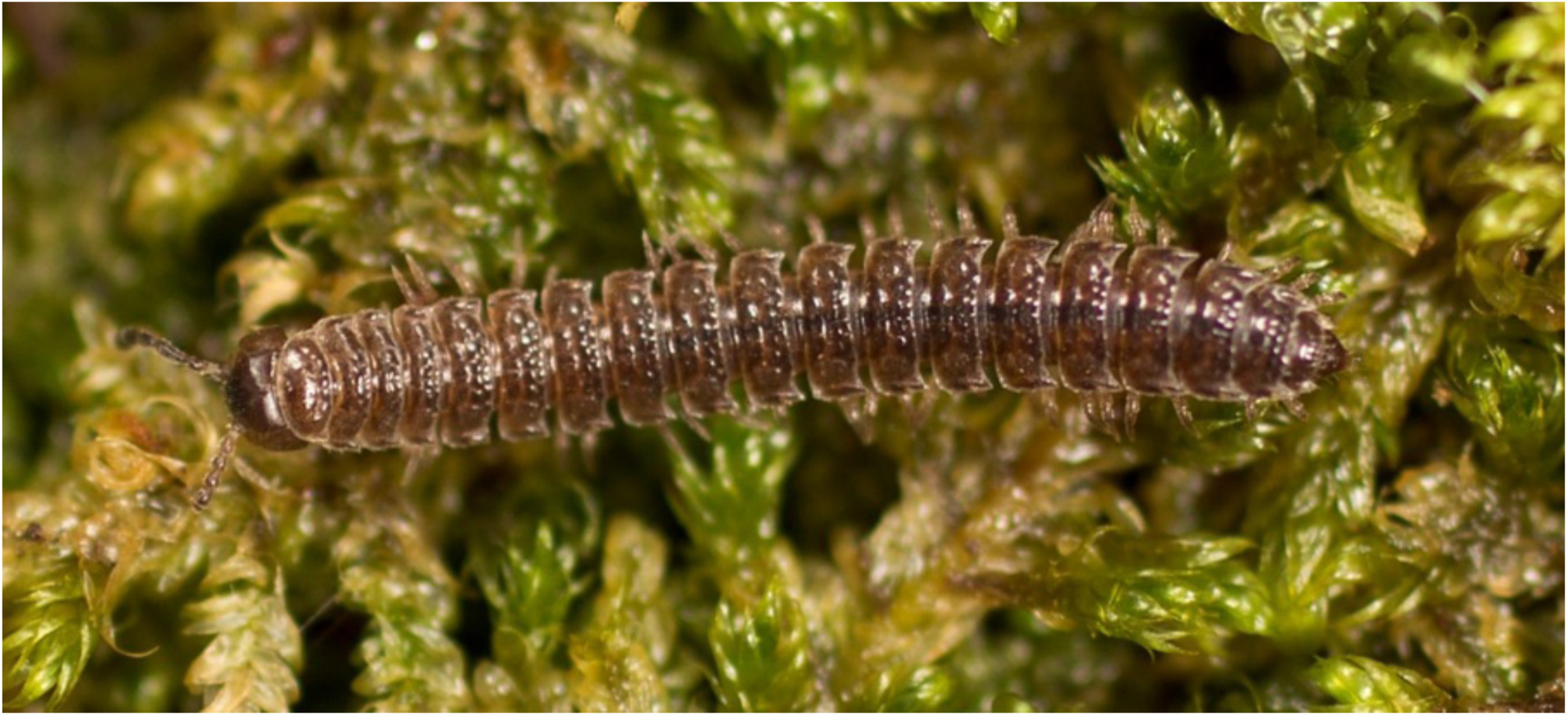
*Pseudopolydesmus paludicolus*, dorsal view of adult female from Colleton County, South Carolina (MPE03525).

**Distribution:** *Pseudopolydesmus paludicolus* is known from the Atlantic Coastal Plain in Virginia and South Carolina (Fig. 14). It has not yet been recorded from North Carolina, but with further collection in swampy habitats, will likely be found to occur in that state.

**Ecology**: A wetland-adapted species, *P. paludicolus* is found exclusively in swamps, often close to the water’s edge. It has been recorded from swamp forests composed of bald cypress (*Taxodium distichum*), black gum (*Nyssa sylvatica*), live oak (*Quercus virginiana*), overcup oak (*Quercus lyrata*), loblolly pine (*Pinus taeda*), and sweet gum (*Liquidambar styraciflua*). It can be found in leaf litter or under logs and other woody debris, and has also been collected within sand dune habitats.

**Pseudopolydesmus pinetorum (Bollman, 1888)**

*Polydesmus americanus* Carl, 1902

*Polydesmus natchitoches* Chamberlin, 1942

*Polydesmus paroicus* Chamberlin, 1942

*Polydesmus hubrichti* Chamberlin, 1943

*Polydesmus modocus* Chamberlin, 1943

Vernacular name: “The Pinelands Flatback”

**Figures 22-24.**

**Diagnosis**: Pulvillus large, rounded distally (Fig. 22). Process *e1* absent, not present as in *P. erasus*; process *e2* medium-sized, fin-shaped, lateral in position, not small and triangular as in *P. minor*; process *e3* large, fin-shaped, and recurved, not absent as in *P. serratus*; process *e4* small, triangular, not absent as in *P. caddo*; process *m1* small, quadrate, not large as in *P. erasus*; process *m2* medium-sized, triangular, not large as in *P. minor*; process *m3* absent, not present as in *P. canadensis*; process *m4* medium-sized, triangular, not absent as in *P. serratus*.

**Figure 22.**
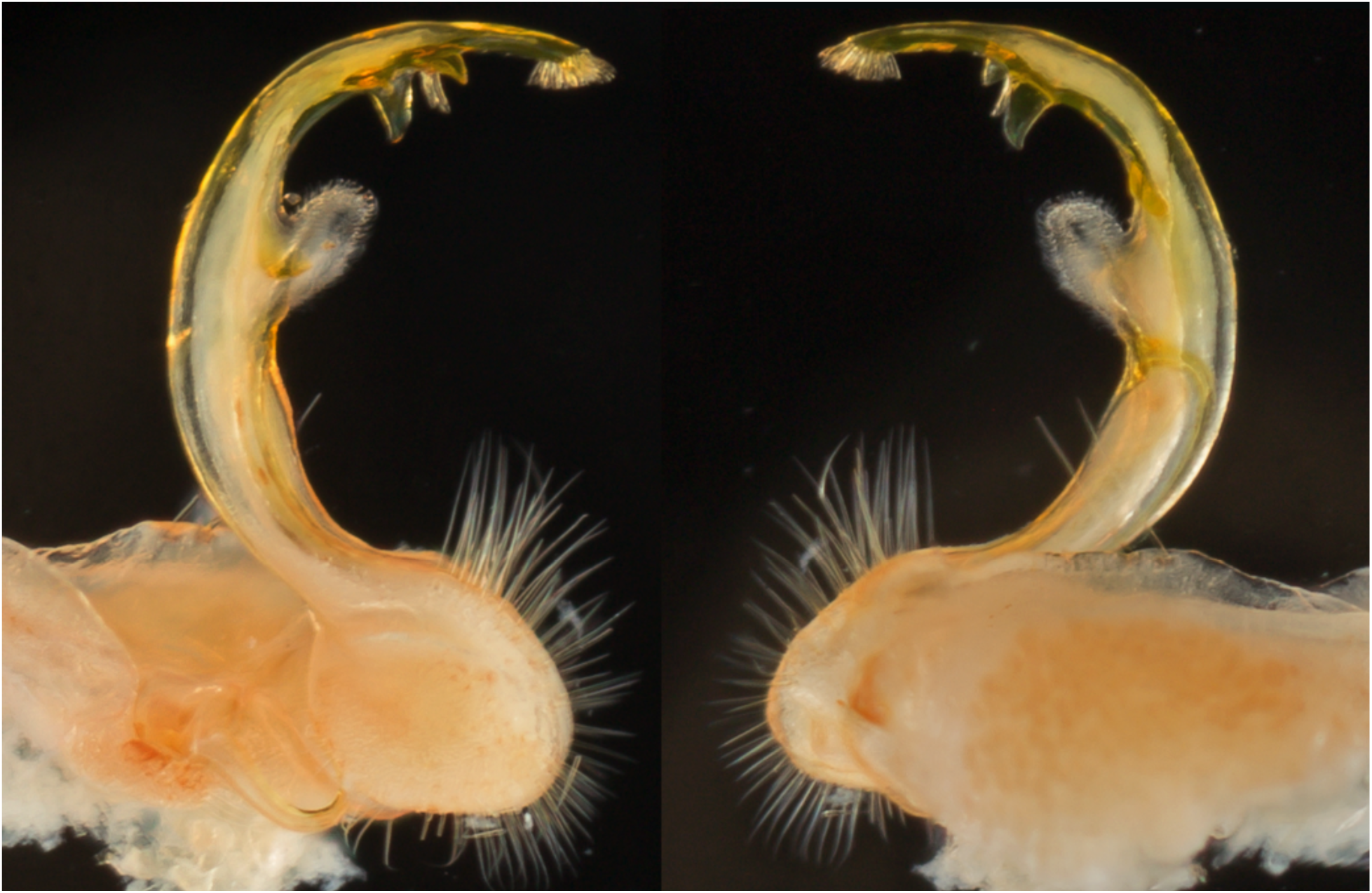
Left gonopod of *Pseudopolydesmus pinetorum* (MPE01006, Newton County, Arkansas), medial view (left) and ectal view (right).

**Color**: Tergites maroon-brown, with paranota light pink to tan in color (Fig. 23). Quadrate paranota often times only slightly pinker than tergites.

**Figure 23.**
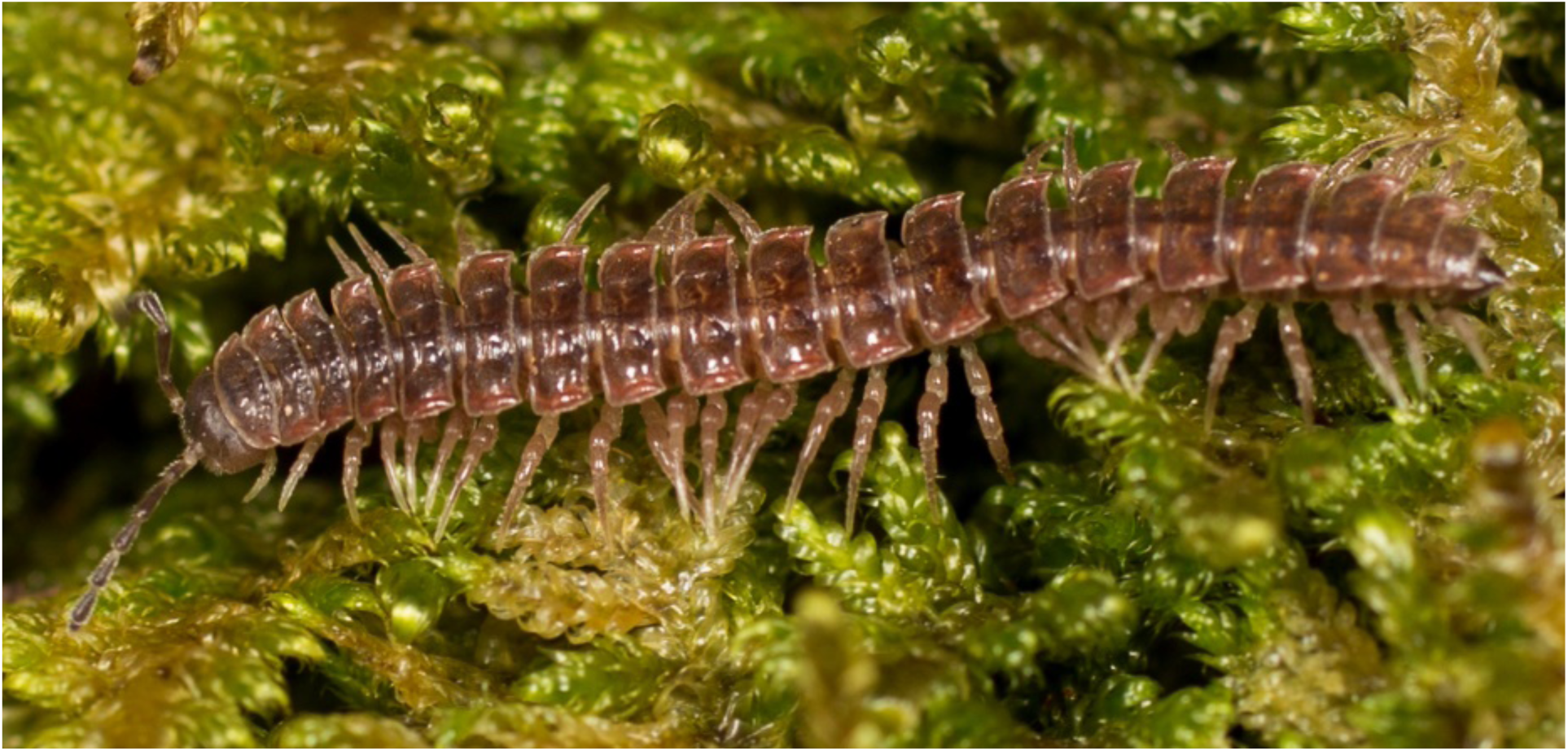
*Pseudopolydesmus pinetorum*, dorsal view of adult male from St. Charles County, Missouri (MPE02753).

**Distribution**: *Pseudopolydesmus pinetorum* is the most common species west of the Mississippi River, but scattered records indicate its presence in farther eastern states (Fig. 24).

**Figure 24.**
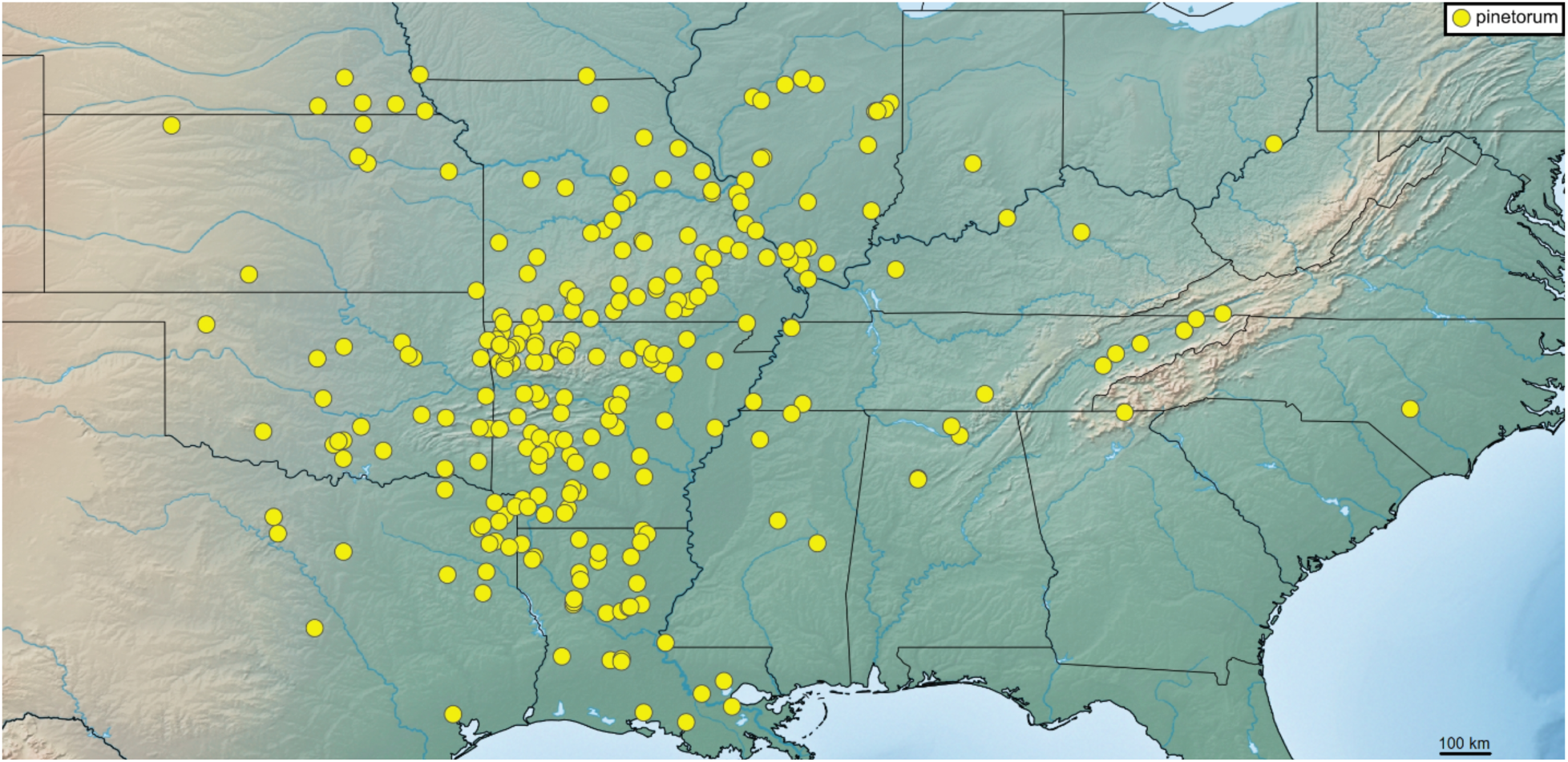
Distribution of *Pseudopolydesmus pinetorum*.

**Ecology**: This species can be found in generalized deciduous woods, as well as in pine forests throughout its range, where it occupies low to middle elevations up to 660 m.

**Pseudopolydesmus serratus (Say, 1821)**

*Polydesmus pennsylvanicus* C. L. Koch, 1847

*Polydesmus scopus* Chamberlin, 1942

*Polydesmus planicolens* Chamberlin, 1942

Vernacular name: “The Common Pink Flatback”

**Figs. 25-27.**

**Diagnosis**: Pulvillus large, slightly rounded distally (Fig. 25). Process *e1* absent, not present as in *P. canadensis*; process *e2* large, triangular, connected to process *m2* by transverse ridge, not connected to another process by a basal stalk as in *P. canadensis*; process *e3* absent, not present as in *P. erasus*; process *e4* small, triangular, not large as in *P. paludicolus*; process *m1* medium-sized, triangular, not large as in *P. minor*; process *m2* large, triangular, connected to process *e2* by transverse ridge, not connected to process *m3* as in *P. paludicolus*; process *m3* absent, not present as in *P. canadensis*; process *m4* absent, not present as in *P. canadensis*.

**Figure 25.**
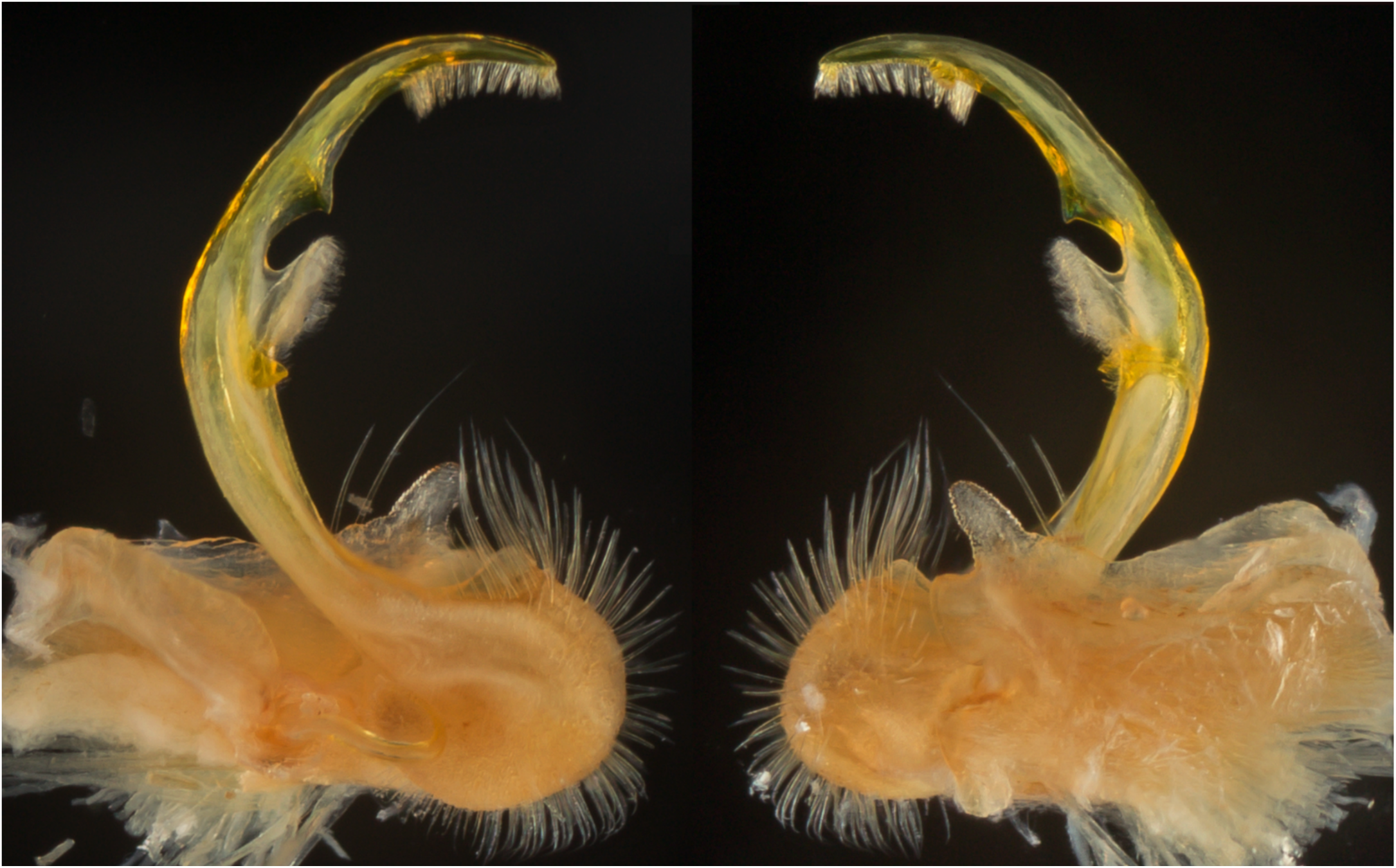
Left gonopod of *Pseudopolydesmus serratus* (MPE00803, Craig County, Virginia), medial view (left) and ectal view (right).

**Color**: Tergites dark brown to light pink, with pink to red paranota (Fig. 26).

**Figure 26.**
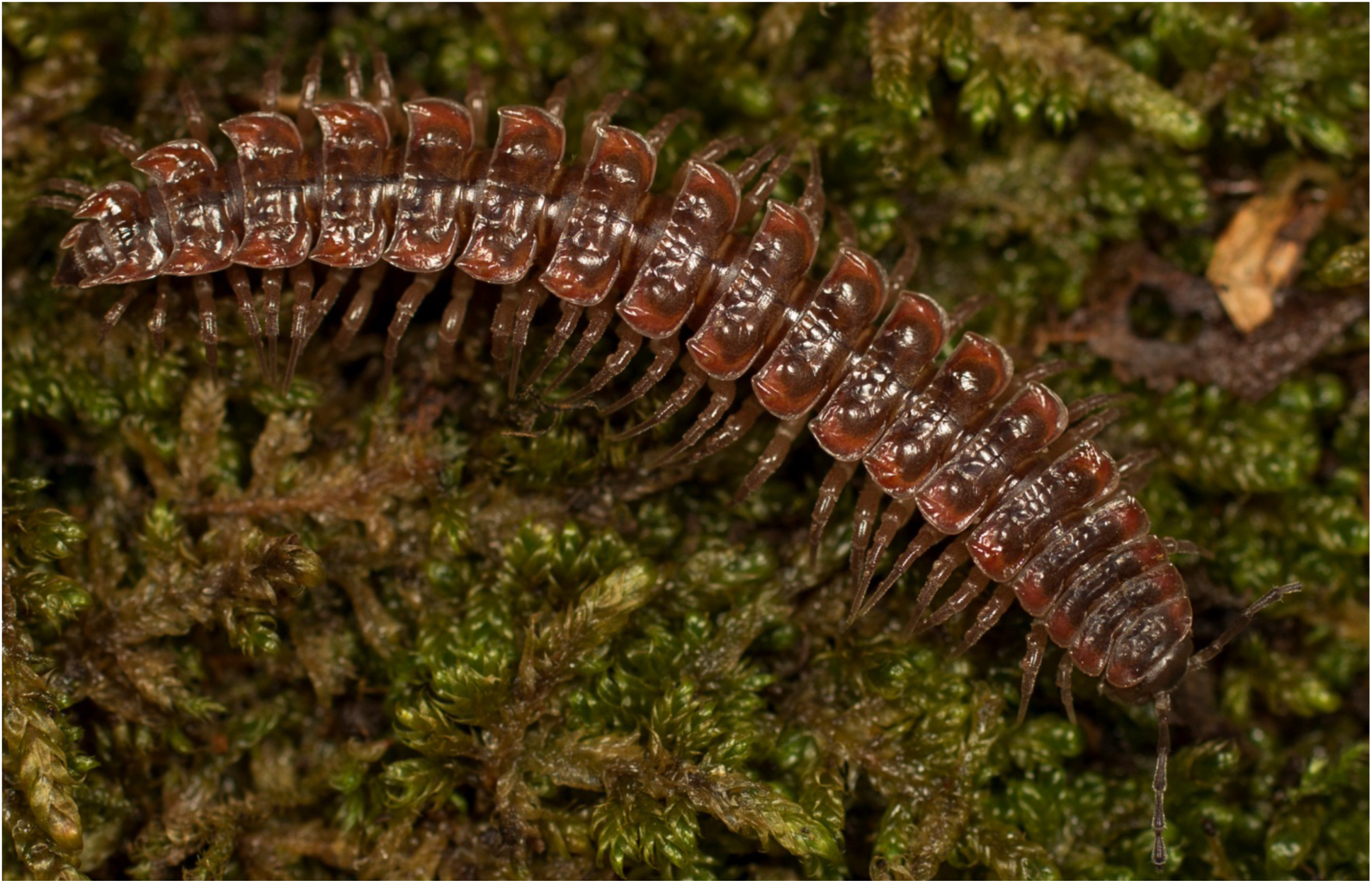
*Pseudopolydesmus serratus,* dorsal view of adult male from Powell County, Kentucky (MPE03105).

**Distribution**: The most widely-distributed species in the genus, *P. serratus* is known from Ontario and Québec south to Florida and Texas (Fig. 27). It is sympatric with every other species in various portions of its large range, but is not as common in the southeastern Coastal Plain and portions of its far western range.

**Figure 27.**
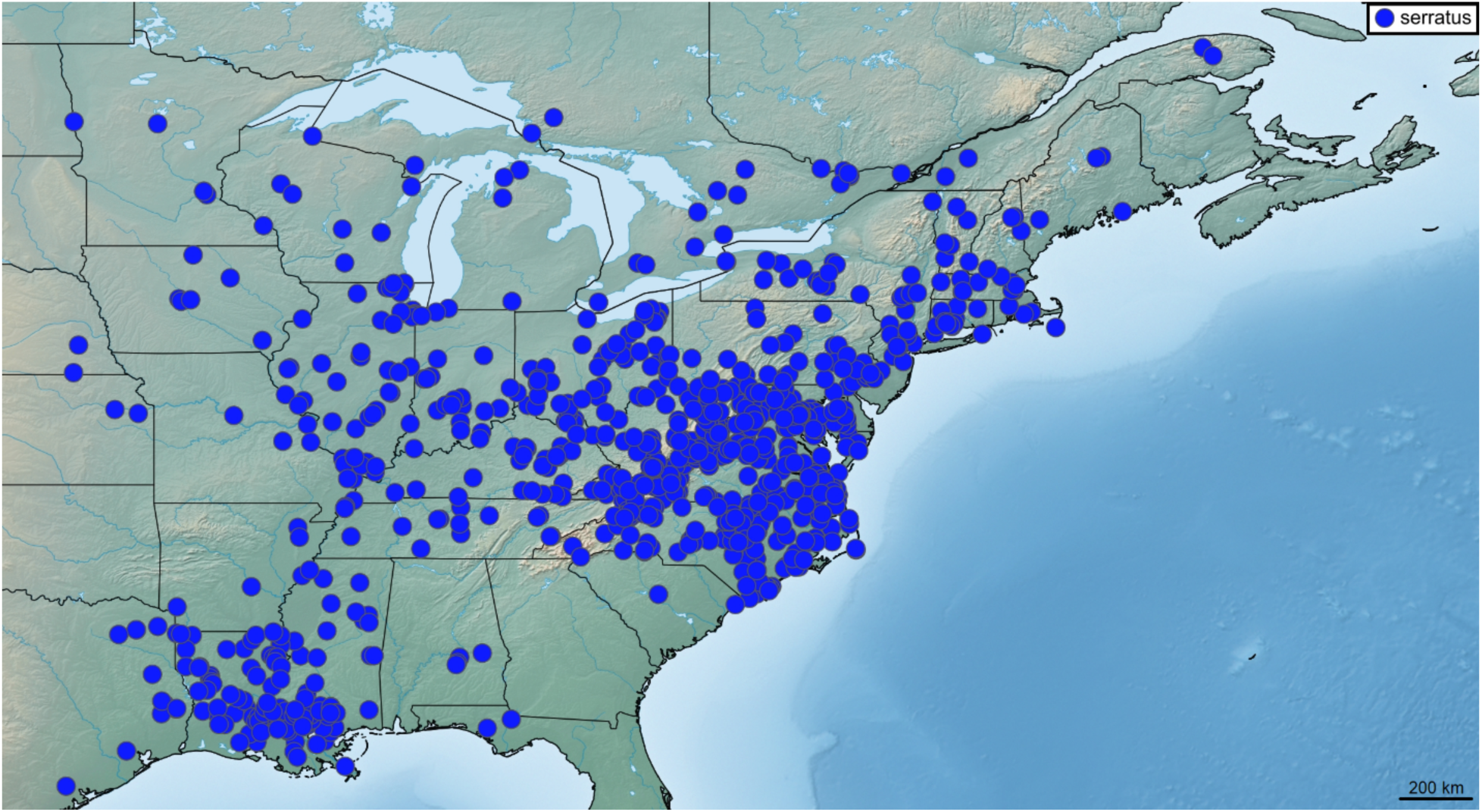
Distribution of *Pseudopolydesmus serratus*.

**Ecology**: A generalist, *Pseudopolydesmus serratus* can be found in almost any suitable millipede habitat within its distribution, though it does not tend to occur at high elevations.

## Discussion

Compared to the morphologically heterogeneous and ill-defined European and Asian polydesmid genera, *Pseudopolydesmus* has been generally assumed to be a monophyletic assemblage, and this is supported by the genetic analyses presented here. The most closely-related genus appears to be *Epanerchodus* (Fig. 3); however, only one of nine other North American polydesmid genera was included, *Scytonotus*. The current concept of eight species in the genus advanced by Sierwald et al. (2019) was supported by the RAxML-inferred phylogeny. Specifically, in each of the mitochondrial gene trees (CO1, 16S) and one of the nuclear gene trees (Enolase), the eight species are recovered as monophyletic. In the remaining two nuclear gene trees (28S, Ef1a), the species were not all monophyletic. In the Ef1a gene tree, six of the species were monophyletic, while *P. canadensis* and *P. collinus* were grouped together in a weakly-supported clade (bootstrap support = 29). In the 28S gene tree, *P. caddo*, *P. minor*, *P. paludicolus, P. pinetorum*, and *P. serratus* were monophyletic, while *P. canadensis*, *P. collinus*, and *P. erasus* were polyphyletic. *Pseudopolydesmus* specimens exhibited very low 28S sequence variation, with 64% of the sequences being identical, along with low clade support (generally < 70, except for *P. caddo* and *P. minor*).

Specimens of the potential new species identified in Hoffman’s notes were recently collected from Russell County, Virginia, for this study. In his notes, Hoffman used the designation “Adocetus” (from the Greek *adoketos*: unexpected) for specimens collected in Russell, Scott, and Washington counties in southwest Virginia, that have the same gonopod processes as in *Pseudopolydesmus serratus*. These individuals have a greatly-curving acropodite, rather than the slightly-curving acropodite typically observed in *P. serratus*. These “Adocetus” specimens were included in the phylogenetic analysis to test if this feature was a distinctive characteristic of a cryptic species, as Hoffman surmised. The results from the RAxML phylogeny indicated that these “Adocetus” specimens are not representative of a cryptic species, as they fall within the *P*. *serratus* clade and do not form their own separate clade. The “Adocetus” specimens are therefore just an example of intraspecific morphological variation within *P*. *serratus*.

The gonopod morphology of *P. erasus* and *P. tallulanus* (Chamberlin, 1943) is similar, and *P. tallulanus* has traditionally been separated by the presence of an additional process, *e3*, on the ectal side of the gonopod; however, specimens conforming to the *P. tallulanus* morphotype nested within the *P. erasus* clade, supporting the synonymy of *P. tallulanus* within *P. erasus* by Hoffman (1999). The gonopod processes of each species are the same, except for the presence or absence of *e3*. Specimens of the *P. tallulanus* morphotype tended to be found at higher elevations in the southern Appalachian Mountains in North Carolina and Georgia, while specimens conforming to the *P. erasus* morphotype tended to be found at lower elevations from Illinois south through Alabama and Georgia, though this did not hold true for every specimen.

Specimens from throughout the distribution of *P. erasus* and that were represented in the molecular phylogeny were examined for the presence of process *e3*, and though many lacked this process entirely, there was some variation. A specimen from Marshall County, Tennessee (MPE01288) was observed to have a very small process *e3*, as did a specimen from Towns County, Georgia (MPE03315), while specimens from Walker County, Georgia (MPE01957) and Tishomingo County, Mississippi (MPE02698) lacked this process entirely. Another specimen from Towns County, Georgia (MPE04338) instead had a very large process *e3*. By examining the morphological variation of these specimens within a molecular phylogenetic framework, none of this variation was found to be restricted to separate clades within *P. erasus*, supporting the conclusion that the *P. tallulanus* morphotype specimens are within the standard morphological variation of *P. erasus*.

Aside from this difference, the traditionally-delineated species of *Pseudopolydesmus,* based solely on gonopod morphology, agree with the genetic analysis. This finding offers great hope for future systematic work based on genetics within the Polydesmidae (and possibly the Polydesmidea as well), offering a more objective basis to base species hypotheses on, instead of relying mainly on small gonopod morphological differences to delimit species and subspecies.

The multivariate morphological analyses were not as clarifying for species delimitation in the genus, however. Body length clearly drove the results of the analysis, with 96% of the variation coming from this component, while the second component accounted for about 2% of the variation. The PCA plot contained six clusters (Fig. 5), and the variables are highly positively correlated, indicating the extent to which they are driven by body size (length). Due to the comparatively uniform somatic morphology in the genus, the modified PCA was not as useful as it might be in a group of taxa that shows more morphological and/or ecological variation. In taxa like *Pseudopolydesmus*, species delimitation based on the genital morphology and genetics, rather than somatic characters, is likely to be a more practical approach.

The previous hypothesis about species relationships within *Pseudopolydesmus* (Withrow 1988) posited two groups: a *canadensis* group (five species) and a *serratus* group (four species), based on the gonopod morphology. This hypothesis was not supported by the species phylogeny inferred by RAxML (Fig. 3) or by the *BEAST species tree (Fig. 6). The phylogenetic analyses each show two clades within *Pseudopolydesmuş* but with varying species compositions. The RAxML tree shows a clade with *P. paludicolus* and *P. serratus* as sister species, with the rest of the species in their own clade (see Results section). In the *BEAST tree, the two major clades within *Pseudopolydesmus* consist of *P. caddo* and *P. minor* as highly-supported sister species, and allied with *P. pinetorum*, while the rest of the species make up the other clade. The RAxML phylogeny has better support values than the *BEAST tree, lending it more credence, but nonetheless, both phylogenies do not support the previous morphological-based hypotheses of species relationships.

These results show biogeographical and ecological patterns within the genus, particularly with regards to adaptation and convergent evolution in wetland habitats. *Pseudopolydesmus paludicolus* was not found to be closely related to *P. minor* and *P. caddo*, the two other wetland-adapted species. This suggests that small body size and adaptation to wetland habitats may have evolved independently twice within the genus. Body length in the three wetland-inhabiting species ranged from 12 – 17 mm, compared to 14 – 36 mm in the rest of the species (Supplementary Material B). All the other species in the genus are general forest species, with some living at higher elevations. Some specimens within *Pseudopolydesmus canadensis* are clustered in a small clade, and all of these specimens are from the southeastern Coastal Plain. However, while they are smaller in body size and have different coloration, they do not appear to be a separate species from *P. canadensis*, though Coastal Plain *P. canadensis* have been thought to warrant subspecific status by previous workers (Hoffman, 1999).

As a result of this study, *Pseudopolydesmus* is now known from 37 U.S. states and two Canadian provinces in central and eastern North America. It is the most widespread polydesmid genus in North America, and the best-studied polydesmid in North America, with summarized information on its morphology (Sierwald et al., 2019; Zahnle et al., 2020) and genetics available. As demonstrated here, future studies on polydesmids will be well-served by incorporating a genetic component for species delimitation.

## Acknowledgements

This research was funded by National Science Foundation grant DEB 2433355 to P. Marek. Thanks to Bronwyn Williams of the North Carolina Museum of Natural Sciences, Kaloyan Ivanov of the Virginia Museum of Natural History, and Thomas McElrath of the Illinois Natural History Survey for facilitating loans of millipede specimens. We thank Maddie Hellier and Sarah Turner for measuring the specimens used in the principal components analysis. We also thank all the collectors who provided specimens, including Sergei Golovatch, Curt Harden, Matt Kasson, Cedric Lee, Wendy Moore, David Kavanaugh, Pavel Nefediev, and Pat Wooden. We thank Jackson Means and Victoria Wong for their assistance in field collection.

Supplemental funding for research was by a National Science Foundation DEB# 1916368 and 1655635, a Theodore Roosevelt Memorial Grant from the American Museum of Natural History, and a Graduate Research Development Program award from the Virginia Tech Graduate Student Assembly. Scientific collection permits were obtained from the relevant organizations and agencies: Cincinnati Museum Center, Edge of Appalachia Preserve; Indiana University, Griffy Woods Preserve; Maryland Department of Natural Resources, Project# 2016DNR119; Maryland State Parks; Georgia State Parks, permit #132016, 142018; Kentucky Department of Parks, permit #1708; University of Illinois, Allerton Park; Champaign County Forest Preserve District; Vermilion County Conservation District; Kentucky State Nature Preserves Commission, permit #2017-09-05; Lexington-Fayette Urban County Government, Raven Run Nature Sanctuary; North Carolina Department of Natural and Cultural Resources, permit #2018_0234; Duke University, Office of Duke Forest, project# R1516-413; South Carolina Department of Parks, Recreation & Tourism, State Park Service, permit# N-26-17; Tennessee Wildlife Resources Agency permit #3914, 1702; Tennessee Department of Environment and Conservation, permit #2017-008; West Virginia Division of Natural Resources, permit #2016.238; Virginia Department of Game and Inland Fisheries, permit #056958; New York State Parks, permit# 2017-MP15-018; Pennsylvania State Parks, permit# 2017-64; Pennsylvania Department of Conservation and Natural Resources, Bureau of Forestry, permit# SFRA-1724; National Parks Service, permit #GRSM-2016-SCI-2369, GRSM-2019-SCI-2369, SHEN-2016-SCI-0018, BLRI-2014-SCI-0033.

