## Supplementary Material A for "Integrative taxonomy of the North American pink flat-backed millipedes (Polydesmidae: *Pseudopolydesmus* Attems, 1898)"

**Appendix X.** List of taxa used in molecular analyses, organized alphabetically by family, genus, and species. Locality refers to country and state/province for localities outside USA. Within USA, locality refers to state and county. Acc# refer to the NCBI Genbank database. All specimens are deposited in the Virginia Tech Insect Collection, Blacksburg, Virginia, USA.

| **Taxon** | **Spc#** | **Latitude** | **Longitude** | **Locality** | **Acc# 16S** | **Acc# EF1a** | **Acc# 28S** | **Acc# COI** | **Acc# Enolase** |
| --- | --- | --- | --- | --- | --- | --- | --- | --- | --- |
| **Cryptodesmidae**  *Kiusiunum sp.* | MPE04845 | 32.802 | 130.2923 | Japan, Nagasaki | - | - | **MT753433** | **MT740080** | - |
| *Peridontodesmus sp.* | MPE04654 | 18.5868 | -95.0762 | Mexico, Veracruz | - | - | **MT753422** | **MT740075** | - |
| **Dalodesmidae**  *Dalodesmidae sp.* | MPE04979 | -24.6388 | 46.6991 | Madagascar, Anosy | - | **MT756837** | - | **MT740087** | - |
| **Haplodesmidae**  *Eutrichodesmus armatocaudatus* | MPE04248 | 20.2934 | 105.6685 | Vietnam, Ninh Bình | - | **MT756815** | **MT753403** | - | - |
| *Eutrichodesmus peculiaris* | MPE04934 | 36.4304 | 140.391 | Japan, Ibaraki | - | **MT756836** | **MT753440** | - | - |
| **Macrosternodesmidae**  *Chaetaspis ohionis* | MPE02995 | 39.4211 | -81.361 | Ohio, Washington | **MT754084** | **MT756758** | **MT753329** | **MT739979** | - |
| *Harpogonopus confluentus* | MPE04221 | 34.037 | -118.734 | California, Los Angeles | - | - | **MT753402** | **MT740056** | - |
| *Kepolydesmus anderisus* | MPE00959 | 47.0847 | -116.115 | Idaho, Shosone | **MT753963** | - | **MT753191** | **MT739846** | - |
| *Nearctodesmus salix* | MPE00982 | 37.8242 | -122.175 | California, Alameda | **MT753968** | - | **MT753198** | **MT739853** | - |
| **Opisotretidae**  *Corypholophus* sp. | MPE04249 | 20.359 | 105.5788 | Vietnam, Ninh Bình | **MT754154** | **MT756816** | **MT753404** | **MT740057** | - |
| Opisotretidae sp. | MPE04933 | 36.4304 | 140.391 | Japan, Ibaraki | - | **MT756835** | **MT753439** | **MT740086** | - |
| **Polydesmidae**  *Brachydesmus furcatus* | MPE04750 | 44.9057 | 37.9764 | Russia, Krasnodar | **MT754171** | - | **MT753426** | **MN232237** | - |
| *Epanerchodus koreanus* | MPE04764 | 43.0833 | 131.4999 | Russia, Primorsky Krai | - | - | **MT753427** | **MN232238** | **MT799659** |
| *Epanerchodus kunashiricus* | MPE03455 | 44.0818 | 145.8747 | Russia, Sakhalin Oblast | - | **MT756784** | **MT753361** | **MN232234** | - |
| *Epanerchodus lacteus* | MPE04884 | 36.4304 | 140.391 | Japan, Ibaraki | - | - | **MT753435** | **MT740082** | - |
| *Epanerchodus polymorphus* | MPE04778 | 43.1884 | 131.9278 | Russia, Primorsky Krai | - | **MT756829** | **MT753428** | **MN232240** | - |
| *Epanerchodus polymorphus* | MPE04779 | 43.0583 | 131.8065 | Russia, Primorsky Krai | - | **MT756830** | **MT753429** | **MN232239** | - |
| *Epanerchodus* sp. | MPE01958 | 27.2052 | 98.6959 | China, Yunnan | **MT754034** | **MT756717** | **MT753271** | **MT739921** | - |
| *Epanerchodus* sp. | MPE01959 | 27.7166 | 98.5032 | China, Yunnan | **MT754035** | **MT756718** | **MT753272** | **MT739922** | - |
| *Epanerchodus* sp. | MPE04847 | 36.1646 | 140.1312 | Japan, Ibaraki | - | **MT756832** | **MT753434** | **MT740081** | - |
| *Epanerchodus* sp. | MPE04897 | 43.0521 | 141.3139 | Japan, Hokkaido | **MT754174** | **MT756833** | **MT753436** | **MT740083** | - |
| *Epanerchodus* sp. | MPE04898 | 43.0521 | 141.3139 | Japan, Hokkaido | - | - | **MT753437** | **MT740084** | - |
| Polydesmidae sp. | MPE04912 | 32.7119 | 130.2165 | Japan, Nagasaki | **MT754175** | **MT756834** | **MT753438** | **MT740085** | - |
| *Polydesmus angustus* | MPE03837 | 37.2125 | -80.4053 | Virginia, Montgomery | **MT754134** | - | **MT753382** | **MT740033** | - |
| *Polydesmus inconstans* | MPE03014 | 53.53 | -113.523 | Canada, Alberta | **MT754087** | - | **MT753332** | **MT739982** | - |
| *Polydesmus vietnamicus* | MPE03839 | 21.4593 | 105.6479 | Vietnam, Vĩnh Phúc | **MT754135** | **MT756799** | **MT753383** | **MT740034** | - |
| *Pseudopolydesmus caddo* | MPE03746 | 30.4554 | -91.827 | Louisiana, Landry | **MT754132** | **MT756797** | **MT753380** | **MT740031** | - |
| *Pseudopolydesmus caddo* | MPE03747 | 30.4554 | -91.827 | Louisiana, Landry | **MT754133** | **MT756798** | **MT753381** | **MT740032** | - |
| *Pseudopolydesmus canadensis* | MPE00024 | 37.0714 | -81.3067 | Virginia, Bland | **MT753928** | **MT756633** | **MT753156** | **MT739814** | **MT799555** |
| *Pseudopolydesmus canadensis* | MPE00318 | 36.1386 | -81.6694 | North Carolina, Watauga | **MT753934** | **MT756639** | **MT753162** | **MT739819** | **MT799558** |
| *Pseudopolydesmus canadensis* | MPE00368 | 37.2764 | -80.4837 | Virginia, Montgomery | **MT753935** | **MT756640** | **MT753163** | **MT739820** | - |
| *Pseudopolydesmus canadensis* | MPE00395 | 37.1018 | -80.8867 | Virginia, Pulaski | **MT753936** | **MT756641** | **MT753164** | **MT739821** | **MT799559** |
| *Pseudopolydesmus canadensis* | MPE00415 | 37.0681 | -80.873 | Virginia, Pulaski | **MT753937** | **MT756642** | **MT753165** | **MT739822** | **MT799560** |
| *Pseudopolydesmus canadensis* | MPE00799 | 37.4035 | -80.3414 | Virginia, Craig | **MT753938** | **MT756643** | **MT753166** | **MT739823** | **MT799561** |
| *Pseudopolydesmus canadensis* | MPE00815 | 35.3554 | -83.9336 | North Carolina, Graham | **MT753941** | **MT756645** | **MT753169** | **MT739826** | **MT799563** |
| *Pseudopolydesmus canadensis* | MPE00841 | 36.7848 | -80.3733 | Virginia, Patrick | **MT753943** | - | **MT753171** | **MT739827** | **MT799565** |
| *Pseudopolydesmus canadensis* | MPE00894 | 37.35 | -80.5835 | Virginia, Giles | **MT753949** | **MT756650** | **MT753177** | **MT739833** | - |
| *Pseudopolydesmus canadensis* | MPE00902 | 36.7848 | -80.3733 | Virginia, Patrick | **MT753951** | **MT756652** | **MT753179** | **MT739835** | **MT799571** |
| *Pseudopolydesmus canadensis* | MPE00903 | 36.7724 | -80.4054 | Virginia, Floyd | **MT753952** | - | **MT753180** | **MT739836** | - |
| *Pseudopolydesmus canadensis* | MPE00905 | 37.2506 | -80.461 | Virginia, Montgomery | **MT753954** | **MT756654** | **MT753182** | **MT739838** | **MT799572** |
| *Pseudopolydesmus canadensis* | MPE00923 | 37.1732 | -81.154 | Virginia, Bland | **MT753955** | **MT756655** | **MT753183** | **MT739839** | **MT799573** |
| *Pseudopolydesmus canadensis* | MPE00929 | 36.9533 | -82.0524 | Virginia, Russell | **MT753956** | **MT756656** | **MT753184** | **MT739840** | **MT799574** |
| *Pseudopolydesmus canadensis* | MPE00946 | 33.7434 | -80.7346 | South Carolina, Calhoun | **MT753960** | **MT756659** | **MT753188** | **MT739844** | - |
| *Pseudopolydesmus canadensis* | MPE00955 | 37.3272 | -80.2426 | Virginia, Roanoke | **MT753962** | **MT756660** | **MT753190** | **MT739845** | **MT799578** |
| *Pseudopolydesmus canadensis* | MPE00986 | 37.0093 | -81.1674 | Virginia, Wythe | **MT753969** | **MT756665** | **MT753199** | **MT739854** | - |
| *Pseudopolydesmus canadensis* | MPE01001 | 36.8176 | -81.4385 | Virginia, Smyth | **MT753971** | **MT756666** | **MT753201** | - | **MT799582** |
| *Pseudopolydesmus canadensis* | MPE01025 | 35.6927 | -83.4799 | Tennessee, Sevier | **MT753975** | **MT756669** | **MT753205** | **MT739858** | - |
| *Pseudopolydesmus canadensis* | MPE01054 | 35.6698 | -82.3628 | North Carolina, Buncombe | **MT753976** | **MT756670** | **MT753206** | **MT739859** | - |
| *Pseudopolydesmus canadensis* | MPE01176 | 35.7528 | -81.3416 | North Carolina, Catawba | **MT753985** | - | **MT753215** | **MT739866** | - |
| *Pseudopolydesmus canadensis* | MPE01276 | 35.9531 | -83.6865 | Tennessee, Knox | **MT753992** | **MT756682** | **MT753223** | **MT739874** | - |
| *Pseudopolydesmus canadensis* | MPE01477 | 35.636 | -83.4937 | Tennessee, Sevier | - | - | **MT753231** | **MT739882** | - |
| *Pseudopolydesmus canadensis* | MPE01485 | 35.4938 | -83.1557 | North Carolina, Jackson | - | - | **MT753232** | **MT739883** | - |
| *Pseudopolydesmus canadensis* | MPE01535 | 36.6866 | -80.4415 | Virginia, Patrick | **MT753999** | **MT756691** | - | **MT739886** | - |
| *Pseudopolydesmus canadensis* | MPE01544 | 35.2147 | -81.8575 | North Carolina, Rutherford | **MT754000** | **MT756692** | **MT753235** | **MT739887** | - |
| *Pseudopolydesmus canadensis* | MPE01650 | 36.9477 | -81.8242 | Virginia, Russell | **MT754004** | **MT756694** | **MT753239** | **MT739891** | **MT799588** |
| *Pseudopolydesmus canadensis* | MPE01682 | 36.9384 | -81.8118 | Virginia, Smythe | **MT754007** | **MT756697** | **MT753242** | **MT739893** | - |
| *Pseudopolydesmus canadensis* | MPE01687 | 37.4647 | -81.0623 | West Virginia, Mercer | **MT754008** | **MT756698** | **MT753243** | **MT739894** | **MT799589** |
| *Pseudopolydesmus canadensis* | MPE01835 | 37.8685 | -79.8713 | Virginia, Alleghany | **MT754017** | - | **MT753252** | **MT739902** | - |
| *Pseudopolydesmus canadensis* | MPE01862 | 38.5948 | -79.1982 | West Virginia, Pendleton | - | **MT756705** | **MT753255** | **MT739905** | **MT799594** |
| *Pseudopolydesmus canadensis* | MPE01865 | 39.6179 | -79.1476 | Maryland, Garrett | **MT754021** | **MT756707** | **MT753257** | **MT739907** | **MT799596** |
| *Pseudopolydesmus canadensis* | MPE01874 | 39.8572 | -79.6774 | Pennsylvania, Fayette | **MT754022** | **MT756708** | **MT753258** | **MT739908** | **MT799597** |
| *Pseudopolydesmus canadensis* | MPE01890 | 38.3089 | -79.3603 | Virginia, Augusta | **MT754025** | **MT756709** | **MT753261** | **MT739911** | **MT799599** |
| *Pseudopolydesmus canadensis* | MPE01893 | 38.33 | -79.2357 | Virginia, Augusta | - | **MT756710** | **MT753263** | **MT739913** | **MT799600** |
| *Pseudopolydesmus canadensis* | MPE01895 | 38.3605 | -79.2048 | Virginia, Augusta | **MT754028** | **MT756712** | **MT753265** | **MT739915** | **MT799601** |
| *Pseudopolydesmus canadensis* | MPE01897 | 38.3634 | -79.2653 | Virginia, Augusta | **MT754029** | **MT756713** | **MT753266** | **MT739916** | **MT799602** |
| *Pseudopolydesmus canadensis* | MPE01912 | 38.2127 | -79.2964 | Virginia, Augusta | **MT754030** | **MT756714** | **MT753267** | **MT739917** | - |
| *Pseudopolydesmus canadensis* | MPE01926 | 37.0257 | -81.0901 | Virginia, Bland | **MT754032** | **MT756715** | **MT753269** | **MT739919** | - |
| *Pseudopolydesmus canadensis* | MPE02047 | 37.3091 | -80.2766 | Virginia, Montgomery | **MT754037** | **MT756720** | **MT753274** | **MT739924** | - |
| *Pseudopolydesmus canadensis* | MPE02053 | 37.8367 | -79.0255 | Virginia, Nelson | **MT754038** | **MT756721** | **MT753275** | **MT739925** | **MT799604** |
| *Pseudopolydesmus canadensis* | MPE02130 | 37.4209 | -80.5093 | Virginia, Giles | **MT754041** | **MT756723** | **MT753278** | **MT739928** | **MT799606** |
| *Pseudopolydesmus canadensis* | MPE02183 | 36.1039 | -83.764 | Tennessee, Knox | - | **MT756724** | **MT753279** | **MT739929** | **MT799607** |
| *Pseudopolydesmus canadensis* | MPE02209 | 36.7067 | -82.5006 | Virginia, Scott | **MT754042** | **MT756725** | **MT753280** | **MT739930** | - |
| *Pseudopolydesmus canadensis* | MPE02275 | 37.7547 | -79.185 | Virginia, Amherst | **MT754043** | **MT756726** | **MT753281** | **MT739931** | **MT799608** |
| *Pseudopolydesmus canadensis* | MPE02296 | 36.7219 | -82.2771 | Virginia, Washington | **MT754045** | **MT756728** | **MT753283** | **MT739933** | **MT799610** |
| *Pseudopolydesmus canadensis* | MPE02389 | 30.7498 | -83.5575 | Georgia, Brooks | - | **MT756731** | **MT753286** | **MT739936** | - |
| *Pseudopolydesmus canadensis* | MPE02390 | 31.3925 | -83.5391 | Georgia, Tift | - | **MT756732** | **MT753287** | **MT739937** | **MT799613** |
| *Pseudopolydesmus canadensis* | MPE02393 | 37.8963 | -79.0007 | Virginia, Augusta | **MT754048** | **MT756733** | **MT753288** | **MT739938** | - |
| *Pseudopolydesmus canadensis* | MPE02412 | 37.0113 | -81.186 | Virginia, Wythe | **MT754051** | **MT756735** | **MT753291** | **MT739941** | **MT799615** |
| *Pseudopolydesmus canadensis* | MPE02435 | 37.5766 | -80.2585 | Virginia, Craig | **MT754052** | - | **MT753292** | **MT739942** | - |
| *Pseudopolydesmus canadensis* | MPE02436 | 37.8963 | -79.0007 | Virginia, Augusta | **MT754053** | **MT756736** | **MT753293** | **MT739943** | - |
| *Pseudopolydesmus canadensis* | MPE02464 | 35.8941 | -83.948 | Tennessee, Knox | **MT754056** | **MT756738** | **MT753296** | **MT739946** | - |
| *Pseudopolydesmus canadensis* | MPE02517 | 36.7825 | -80.3997 | Virginia, Floyd | **MT754062** | - | **MT753302** | **MT739952** | **MT799616** |
| *Pseudopolydesmus canadensis* | MPE02521 | 37.4357 | -80.5482 | Virginia, Giles | **MT754063** | **MT756742** | **MT753303** | **MT739953** | **MT799617** |
| *Pseudopolydesmus canadensis* | MPE02531 | 37.4717 | -80.562 | West Virginia, Monroe | **MT754064** | **MT756743** | **MT753304** | **MT739954** | **MT799618** |
| *Pseudopolydesmus canadensis* | MPE02614 | 36.7181 | -80.3246 | Virginia, Patrick | **MT754068** | - | **MT753308** | **MT739958** | **MT799620** |
| *Pseudopolydesmus canadensis* | MPE02636 | 36.8063 | -81.4272 | Virginia, Smyth | **MT754069** | **MT756747** | **MT753309** | **MT739959** | - |
| *Pseudopolydesmus canadensis* | MPE02675 | 37.4667 | -83.9164 | Kentucky, Jackson | **MT754070** | **MT756748** | **MT753310** | **MT739960** | - |
| *Pseudopolydesmus canadensis* | MPE02841 | 38.2301 | -80.2353 | West Virginia, Pocahontas | **MT754079** | **MT756755** | **MT753322** | **MT739972** | - |
| *Pseudopolydesmus canadensis* | MPE02860 | 36.627 | -81.503 | Virginia, Grayson | **MT754082** | - | **MT753325** | **MT739975** | - |
| *Pseudopolydesmus canadensis* | MPE02927 | 35.9025 | -82.5213 | Tennessee, Madison | **MT754083** | - | **MT753328** | **MT739978** | **MT799629** |
| *Pseudopolydesmus canadensis* | MPE03012 | 37.3866 | -81.8729 | West Virginia, McDowell | **MT754086** | **MT756759** | **MT753331** | **MT739981** | - |
| *Pseudopolydesmus canadensis* | MPE03029 | 40.0394 | -78.6471 | Pennsylvania, Bedford | **MT754091** | **MT756763** | - | **MT739986** | - |
| *Pseudopolydesmus canadensis* | MPE03032 | 42.1774 | -75.4228 | New York, Broome | **MT754093** | **MT756765** | **MT753337** | **MT739988** | **MT799631** |
| *Pseudopolydesmus canadensis* | MPE03060 | 42.5433 | -76.6018 | New York, Tompkins | **MT754095** | **MT756767** | **MT753339** | **MT739990** | **MT799633** |
| *Pseudopolydesmus canadensis* | MPE03068 | 41.1034 | -77.2435 | Pennsylvania, Clinton | **MT754097** | **MT756768** | **MT753341** | **MT739992** | - |
| *Pseudopolydesmus canadensis* | MPE03143 | 37.8915 | -84.3972 | Kentucky, Fayette | - | **MT756776** | **MT753349** | **MT740000** | **MT799639** |
| *Pseudopolydesmus canadensis* | MPE03181 | 37.2897 | -82.2999 | Virginia, Dickenson | **MT754104** | - | **MT753350** | **MT740001** | **MT799640** |
| *Pseudopolydesmus canadensis* | MPE03223 | 38.0455 | -83.5494 | Kentucky, Menifee | **MT754106** | - | - | **MT740003** | **MT799641** |
| *Pseudopolydesmus canadensis* | MPE03322 | 35.4315 | -82.2435 | North Carolina, Rutherford | **MT754111** | **MT756780** | **MT753355** | **MT740008** | **MT799642** |
| *Pseudopolydesmus canadensis* | MPE03491 | 37.9021 | -79.5882 | Virginia, Rockbridge | **MT754116** | **MT756785** | **MT753362** | **MT740013** | **MT799644** |
| *Pseudopolydesmus canadensis* | MPE03544 | 33.064 | -80.6164 | South Carolina, Colleton | **MT754121** | **MT756788** | **MT753367** | **MT740018** | **MT799649** |
| *Pseudopolydesmus canadensis* | MPE03553 | 34.0307 | -79.7816 | South Carolina, Florence | **MT754122** | **MT756789** | **MT753369** | **MT740020** | - |
| *Pseudopolydesmus canadensis* | MPE03556 | 33.5122 | -80.49 | South Carolina, Orangeburg | **MT754123** | **MT756790** | **MT753370** | **MT740021** | - |
| *Pseudopolydesmus canadensis* | MPE03558 | 35.2396 | -81.8716 | North Carolina, Rutherford | **MT754124** | - | **MT753371** | **MT740022** | **MT799650** |
| *Pseudopolydesmus canadensis* | MPE03560 | 34.3334 | -79.2683 | South Carolina, Dillon | **MT754125** | **MT756791** | **MT753372** | **MT740023** | **MT799651** |
| *Pseudopolydesmus canadensis* | MPE03562 | 34.1946 | -80.1833 | South Carolina, Lee | - | **MT756792** | **MT753373** | **MT740024** | **MT799652** |
| *Pseudopolydesmus canadensis* | MPE03590 | 34.1715 | -82.3955 | South Carolina, Abbeville | **MT754126** | **MT756793** | **MT753374** | **MT740025** | - |
| *Pseudopolydesmus canadensis* | MPE03593 | 34.139 | -82.315 | South Carolina, Abbeville | **MT754127** | **MT756794** | **MT753375** | **MT740026** | - |
| *Pseudopolydesmus canadensis* | MPE03595 | 35.3491 | -81.851 | North Carolina, Rutherford | **MT754128** | - | **MT753376** | **MT740027** | - |
| *Pseudopolydesmus canadensis* | MPE03664 | 32.5721 | -81.3106 | South Carolina, Hampton | **MT754131** | **MT756796** | **MT753379** | **MT740030** | **MT799653** |
| *Pseudopolydesmus canadensis* | MPE03876 | 34.127 | -82.2767 | South Carolina, Greenwood | **MT754137** | **MT756801** | **MT753385** | **MT740036** | - |
| *Pseudopolydesmus canadensis* | MPE04011 | 38.452 | -79.1144 | Virginia, Rockingham | **MT754147** | **MT756809** | **MT753395** | **MT740048** | **MT799654** |
| *Pseudopolydesmus canadensis* | MPE04042 | 36.5703 | -82.2356 | Tennessee, Sullivan | **MT754149** | **MT756810** | **MT753397** | **MT740050** | - |
| *Pseudopolydesmus canadensis* | MPE04070 | 36.2652 | -82.1235 | Tennessee, Carter | **MT754150** | **MT756811** | - | **MT740051** | - |
| *Pseudopolydesmus canadensis* | MPE04101 | 36.277 | -82.3461 | Tennessee, Washington | **MT754151** | **MT756812** | **MT753398** | **MT740052** | - |
| *Pseudopolydesmus canadensis* | MPE04105 | 36.1742 | -82.2982 | Tennessee, Unicoi | - | **MT756813** | **MT753399** | **MT740053** | - |
| *Pseudopolydesmus canadensis* | MPE04136 | 36.0482 | -82.5615 | Tennessee, Unicoi | **MT754152** | **MT756814** | **MT753400** | **MT740054** | - |
| *Pseudopolydesmus canadensis* | MPE04172 | 35.9609 | -82.8753 | Tennessee, Greene | **MT754153** | - | **MT753401** | **MT740055** | - |
| *Pseudopolydesmus canadensis* | MPE04347 | 35.2851 | -82.8286 | North Carolina, Transylvania | - | **MT756819** | **MT753408** | **MT740061** | - |
| *Pseudopolydesmus canadensis* | MPE04397 | 35.8561 | -82.0876 | North Carolina, Mitchell | **MT754159** | **MT756821** | **MT753410** | **MT740063** | - |
| *Pseudopolydesmus canadensis* | MPE04520 | 35.6136 | -82.2247 | North Carolina, McDowell | **MT754160** | - | **MT753412** | **MT740065** | - |
| *Pseudopolydesmus canadensis* | MPE04527 | 35.4057 | -82.6442 | North Carolina, Henderson | **MT754161** | - | **MT753414** | **MT740067** | - |
| *Pseudopolydesmus canadensis* | MPE04546 | 35.9427 | -83.0866 | South Carolina, Oconee | **MT754163** | **MT756824** | **MT753416** | **MT740069** | - |
| *Pseudopolydesmus canadensis* | MPE04708 | 35.0286 | -77.135 | North Carolina, Jones | **MT754169** | **MT756828** | **MT753424** | **MT740077** | - |
| *Pseudopolydesmus canadensis* | MPE04790 | 38.1024 | -79.9672 | West Virginia, Pocahontas | **MT754173** | **MT756831** | **MT753432** | **MT740079** | **MT799660** |
| *Pseudopolydesmus canadensis* | SPC000222 | 36.4088 | -81.5861 | North Carolina, Ashe | **MT754176** | - | **MT753441** | **MT740088** | - |
| *Pseudopolydesmus canadensis* | SPC001102 | 38.2897 | -80.5353 | West Virginia, Nicholas | **MT754177** | **MT756838** | **MT753442** | **MT740089** | - |
| *Pseudopolydesmus canadensis* | SPC001162 | 35.9648 | -82.9428 | Tennessee, Cocke | **MT754178** | **MT756839** | **MT753443** | **MT740090** | - |
| *Pseudopolydesmus collinus* | MPE00248 | 36.9662 | -80.4179 | Virginia, Floyd | **MT753930** | **MT756635** | **MT753158** | **MT739816** | **MT799557** |
| *Pseudopolydesmus collinus* | MPE00842 | 36.7848 | -80.3733 | Virginia, Patrick | **MT753944** | **MT756647** | **MT753172** | **MT739828** | - |
| *Pseudopolydesmus collinus* | MPE00846 | 36.8706 | -80.4416 | Virginia, Floyd | **MT753945** | - | **MT753173** | **MT739829** | **MT799566** |
| *Pseudopolydesmus collinus* | MPE00971 | 36.8132 | -80.3495 | Virginia, Floyd | **MT753965** | **MT756662** | **MT753194** | **MT739849** | - |
| *Pseudopolydesmus collinus* | MPE00974 | 38.7308 | -79.4912 | West Virginia, Pendleton | **MT753966** | **MT756663** | **MT753195** | **MT739850** | - |
| *Pseudopolydesmus collinus* | MPE01737 | 38.7213 | -83.4335 | Ohio, Adams | **MT754010** | - | **MT753245** | **MT739896** | **MT799591** |
| *Pseudopolydesmus collinus* | MPE01751 | 37.8338 | -81.0735 | West Virginia, Raleigh | **MT754013** | **MT756701** | **MT753248** | **MT739898** | - |
| *Pseudopolydesmus collinus* | MPE01776 | 38.7535 | -85.4222 | Indiana, Jefferson | **MT754014** | **MT756702** | **MT753249** | **MT739899** | - |
| *Pseudopolydesmus collinus* | MPE02494 | 36.7836 | -80.4 | Virginia, Floyd | **MT754060** | **MT756740** | **MT753300** | **MT739950** | - |
| *Pseudopolydesmus collinus* | MPE02498 | 37.1369 | -80.1108 | Virginia, Roanoke | **MT754061** | **MT756741** | **MT753301** | **MT739951** | - |
| *Pseudopolydesmus collinus* | MPE03140 | 38.3738 | -83.1142 | Kentucky, Carter | - | **MT756775** | **MT753348** | **MT739999** | **MT799638** |
| *Pseudopolydesmus collinus* | MPE03440 | 36.55 | -79.35 | Virginia, Pittsylvania | **MT754113** | **MT756782** | **MT753358** | **MT740011** | **MT799643** |
| *Pseudopolydesmus collinus* | MPE03540 | 36.6062 | -80.772 | Virginia, Carroll | **MT754120** | **MT756787** | **MT753366** | **MT740017** | **MT799648** |
| *Pseudopolydesmus erasus* | MPE00295 | 34.2976 | -83.1577 | Georgia, Franklin | **MT753932** | **MT756637** | **MT753160** | **MT739818** | - |
| *Pseudopolydesmus erasus* | MPE00814 | 35.0815 | -83.2359 | North Carolina, Macon | **MT753940** | **MT756644** | **MT753168** | **MT739825** | - |
| *Pseudopolydesmus erasus* | MPE00904 | 35.0815 | -83.2359 | North Carolina, Macon | **MT753953** | **MT756653** | **MT753181** | **MT739837** | - |
| *Pseudopolydesmus erasus* | MPE00933 | 35.3554 | -83.9336 | North Carolina, Graham | **MT753958** | **MT756658** | **MT753186** | **MT739842** | **MT799576** |
| *Pseudopolydesmus erasus* | MPE00943 | 34.9075 | -83.1807 | South Carolina, Oconee | **MT753959** | - | **MT753187** | **MT739843** | - |
| *Pseudopolydesmus erasus* | MPE01229 | 35.6612 | -85.3464 | Tennessee, Van Buren | **MT753988** | **MT756678** | **MT753218** | **MT739869** | **MT799585** |
| *Pseudopolydesmus erasus* | MPE01256 | 35.3695 | -84.9081 | Tennessee, Meigs | **MT753990** | **MT756680** | **MT753220** | **MT739871** | **MT799586** |
| *Pseudopolydesmus erasus* | MPE01270 | 34.5704 | -84.2445 | Georgia, Dawson | **MT753991** | - | **MT753221** | **MT739872** | - |
| *Pseudopolydesmus erasus* | MPE01273 | 34.7438 | -86.5113 | Alabama, Madison | - | **MT756681** | **MT753222** | **MT739873** | - |
| *Pseudopolydesmus erasus* | MPE01288 | 35.5912 | -86.7026 | Tennesee, Marshall | - | **MT756685** | **MT753226** | **MT739877** | **MT799587** |
| *Pseudopolydesmus erasus* | MPE01311 | 34.1755 | -86.8171 | Alabama, Cullman | **MT753994** | **MT756686** | **MT753227** | **MT739878** | - |
| *Pseudopolydesmus erasus* | MPE01326 | 36.1314 | -85.4438 | Tennessee, Putnam | **MT753995** | **MT756687** | **MT753228** | **MT739879** | - |
| *Pseudopolydesmus erasus* | MPE01360 | 34.5731 | -84.1736 | Georgia, Lumpkin | **MT753996** | - | **MT753229** | **MT739880** | - |
| *Pseudopolydesmus erasus* | MPE01486 | 35.4938 | -83.1557 | North Carolina, Jackson | **MT753998** | **MT756689** | **MT753233** | **MT739884** | - |
| *Pseudopolydesmus erasus* | MPE01502 | 35.0288 | -83.2823 | North Carolina, Macon | - | **MT756690** | **MT753234** | **MT739885** | - |
| *Pseudopolydesmus erasus* | MPE01957 | 34.9338 | -85.3715 | Georgia, Walker | **MT754033** | **MT756716** | **MT753270** | **MT739920** | - |
| *Pseudopolydesmus erasus* | MPE02696 | 36.1017 | -87.2853 | Tennessee, Dickson | **MT754072** | - | **MT753312** | **MT739962** | **MT799621** |
| *Pseudopolydesmus erasus* | MPE02698 | 34.8051 | -88.3063 | Mississippi, Tishomingo | **MT754073** | - | **MT753313** | **MT739963** | **MT799622** |
| *Pseudopolydesmus erasus* | MPE02805 | 37.4964 | -89.3572 | Illinois, Union | **MT754078** | **MT756754** | **MT753321** | **MT739971** | **MT799627** |
| *Pseudopolydesmus erasus* | MPE03315 | 34.8364 | -83.771 | Georgia, Towns | **MT754109** | - | **MT753353** | **MT740006** | - |
| *Pseudopolydesmus erasus* | MPE03319 | 34.8408 | -83.8022 | Georgia, Union | **MT754110** | **MT756779** | **MT753354** | **MT740007** | - |
| *Pseudopolydesmus erasus* | MPE04338 | 34.9688 | -83.7733 | Georgia, Towns | **MT754156** | **MT756817** | **MT753406** | **MT740059** | - |
| *Pseudopolydesmus erasus* | MPE04368 | 34.7391 | -83.3887 | Georgia, Rabun | **MT754158** | **MT756820** | **MT753409** | **MT740062** | - |
| *Pseudopolydesmus erasus* | MPE04417 | 34.7701 | -83.9163 | Georgia, Union | - | **MT756822** | **MT753411** | **MT740064** | - |
| *Pseudopolydesmus erasus* | MPE04526 | 35.4057 | -82.6442 | North Carolina, Henderson | - | - | **MT753413** | **MT740066** | **MT799656** |
| *Pseudopolydesmus erasus* | MPE04534 | 34.7262 | -83.7216 | Georgia, White | **MT754162** | **MT756823** | **MT753415** | **MT740068** | **MT799657** |
| *Pseudopolydesmus erasus* | MPE04547 | 35.9427 | -83.0866 | South Carolina, Oconee | **MT754164** | **MT756825** | **MT753417** | **MT740070** | - |
| *Pseudopolydesmus minor* | MPE03910 | 40.0589 | -87.9822 | Illinois, Champaign | **MT754142** | **MT756804** | **MT753390** | **MT740041** | - |
| *Pseudopolydesmus minor* | MPE03953 | 40.0009 | -88.6475 | Illinois, Piatt | **MT754146** | **MT756808** | **MT753394** | **MT740047** | - |
| *Pseudopolydesmus paludicolus* | MPE01167 | 36.567 | -77.4341 | Virginia, Greensville | **MT753982** | **MT756674** | **MT753212** | - | - |
| *Pseudopolydesmus paludicolus* | MPE01170 | 36.567 | -77.4341 | Virginia, Greensville | **MT753983** | **MT756675** | **MT753213** | - | **MT799583** |
| *Pseudopolydesmus paludicolus* | MPE01178 | 36.6927 | -75.9258 | Virginia, Virginia Beach | **MT753986** | **MT756676** | **MT753216** | **MT739867** | - |
| *Pseudopolydesmus paludicolus* | MPE03444 | 36.862 | -77.1876 | Virginia, Sussex | **MT754115** | **MT756783** | **MT753360** | - | - |
| *Pseudopolydesmus paludicolus* | MPE03494 | 33.064 | -80.6164 | South Carolina, Colleton | **MT754117** | **MT756786** | **MT753363** | **MT740014** | **MT799645** |
| *Pseudopolydesmus paludicolus* | MPE03644 | 33.064 | -80.6164 | South Carolina, Colleton | **MT754130** | **MT756795** | **MT753378** | **MT740029** | - |
| *Pseudopolydesmus pinetorum* | MPE01002 | 37.3145 | -90.88 | Missouri, Reynolds | **MT753972** | **MT756667** | **MT753202** | **MT739856** | - |
| *Pseudopolydesmus pinetorum* | MPE01004 | 36.1637 | -94.4374 | Arkansas, Benton | **MT753973** | **MT756668** | **MT753203** | **MT739857** | - |
| *Pseudopolydesmus pinetorum* | MPE01006 | 36.0252 | -93.3522 | Arkansas, Newton | **MT753974** | - | **MT753204** | - | - |
| *Pseudopolydesmus pinetorum* | MPE01070 | 39.1775 | -96.5762 | Kansas, Riley | **MT753977** | **MT756671** | **MT753207** | **MT739860** | - |
| *Pseudopolydesmus pinetorum* | MPE01162 | 34.5211 | -93.3797 | Arkansas, Garland | **MT753980** | **MT756673** | **MT753210** | **MT739863** | - |
| *Pseudopolydesmus pinetorum* | MPE01163 | 34.6222 | -93.8025 | Arkansas, Montgomery | **MT753981** | - | **MT753211** | **MT739864** | - |
| *Pseudopolydesmus pinetorum* | MPE01247 | 36.1301 | -83.5056 | Tennessee, Jefferson | **MT753989** | **MT756679** | **MT753219** | **MT739870** | - |
| *Pseudopolydesmus pinetorum* | MPE01281 | 36.1301 | -83.5056 | Tennessee, Jefferson | - | **MT756683** | **MT753224** | **MT739875** | - |
| *Pseudopolydesmus pinetorum* | MPE01283 | 36.3511 | -82.7674 | Tennessee, Greene | **MT753993** | **MT756684** | **MT753225** | **MT739876** | - |
| *Pseudopolydesmus pinetorum* | MPE02119 | 35.1694 | -93.6452 | Arkansas, Logan | **MT754039** | **MT756722** | **MT753276** | **MT739926** | **MT799605** |
| *Pseudopolydesmus pinetorum* | MPE02715 | 37.1322 | -92.3241 | Missouri, Wright | **MT754075** | - | **MT753315** | **MT739965** | - |
| *Pseudopolydesmus pinetorum* | MPE02718 | 39.1762 | -86.3395 | Indiana, Brown | **MT754076** | **MT756751** | **MT753316** | **MT739966** | **MT799624** |
| *Pseudopolydesmus pinetorum* | MPE02739 | 36.0376 | -93.3412 | Arkansas, Newton | **MT754077** | **MT756752** | **MT753317** | **MT739967** | - |
| *Pseudopolydesmus pinetorum* | MPE02752 | 38.6706 | -90.7515 | Missouri, St. Charles | - | **MT756753** | **MT753318** | **MT739968** | **MT799625** |
| *Pseudopolydesmus pinetorum* | MPE02758 | 36.4307 | -93.7576 | Arkansas, Carroll | - | - | **MT753319** | **MT739969** | - |
| *Pseudopolydesmus pinetorum* | MPE03912 | 40.0589 | -87.9822 | Illinois, Champaign | **MT754143** | **MT756805** | **MT753391** | **MT740042** | - |
| *Pseudopolydesmus pinetorum* | MPE03918 | 40.2033 | -87.7338 | Illinois, Vermilion | - | **MT756806** | **MT753392** | **MT740045** | - |
| *Pseudopolydesmus pinetorum* | MPE04339 | 34.9688 | -83.7733 | Georgia, Towns | **MT754157** | **MT756818** | **MT753407** | **MT740060** | - |
| *Pseudopolydesmus pinetorum* | MPE04597 | 34.7354 | -86.7019 | Alabama, Madison | **MT754165** | **MT756826** | **MT753418** | **MT740071** | - |
| *Pseudopolydesmus pinetorum* | MPE04612 | 36.6398 | -82.1123 | Virginia, Washington | **MT754167** | - | **MT753420** | **MT740073** | - |
| *Pseudopolydesmus serratus* | MPE00143 | 37.5264 | -80.9879 | West Virginia, Mercer | **MT753929** | **MT756634** | **MT753157** | **MT739815** | **MT799556** |
| *Pseudopolydesmus serratus* | MPE00271 | 37.0148 | -81.41 | Virginia, Bland | **MT753931** | **MT756636** | **MT753159** | **MT739817** | - |
| *Pseudopolydesmus serratus* | MPE00299 | 37.9413 | -78.4747 | Virginia, Albemarle | **MT753933** | **MT756638** | **MT753161** | - | - |
| *Pseudopolydesmus serratus* | MPE00803 | 37.4035 | -80.3414 | Virginia, Craig | **MT753939** | - | **MT753167** | **MT739824** | **MT799562** |
| *Pseudopolydesmus serratus* | MPE00816 | 37.2116 | -80.4895 | Virginia, Montgomery | **MT753942** | **MT756646** | **MT753170** | - | **MT799564** |
| *Pseudopolydesmus serratus* | MPE00848 | 36.8132 | -80.3495 | Virginia, Floyd | **MT753946** | - | **MT753174** | **MT739830** | **MT799567** |
| *Pseudopolydesmus serratus* | MPE00889 | 37.3984 | -76.5525 | Virginia, Gloucester | **MT753947** | **MT756648** | **MT753175** | **MT739831** | **MT799568** |
| *Pseudopolydesmus serratus* | MPE00890 | 37.3501 | -80.5833 | Virginia, Giles | **MT753948** | **MT756649** | **MT753176** | **MT739832** | **MT799569** |
| *Pseudopolydesmus serratus* | MPE00898 | 36.0741 | -79.0007 | North Carolina, Orange | **MT753950** | **MT756651** | **MT753178** | **MT739834** | **MT799570** |
| *Pseudopolydesmus serratus* | MPE00930 | 36.9533 | -82.0524 | Virginia, Russell | **MT753957** | **MT756657** | **MT753185** | **MT739841** | **MT799575** |
| *Pseudopolydesmus serratus* | MPE00952 | 37.2939 | -76.6462 | Virginia, York | **MT753961** | **-** | **MT753189** | - | **MT799577** |
| *Pseudopolydesmus serratus* | MPE00962 | 35.7175 | -77.521 | North Carolina, Pitt | - | **MT756661** | **MT753192** | **MT739847** | **MT799579** |
| *Pseudopolydesmus serratus* | MPE00967 | 36.8132 | -80.3495 | Virginia, Floyd | **MT753964** | - | **MT753193** | **MT739848** | **MT799580** |
| *Pseudopolydesmus serratus* | MPE00976 | 38.7308 | -79.4912 | West Virginia, Pendleton | **MT753967** | - | **MT753196** | **MT739851** | - |
| *Pseudopolydesmus serratus* | MPE00988 | 37.0093 | -81.1674 | Virginia, Wythe | **MT753970** | - | **MT753200** | **MT739855** | **MT799581** |
| *Pseudopolydesmus serratus* | MPE01117 | 34.2393 | -77.8228 | North Carolina, New Hanover | **MT753978** | - | **MT753208** | **MT739861** | - |
| *Pseudopolydesmus serratus* | MPE01121 | 35.8366 | -78.7623 | North Carolina, Wake | **MT753979** | **MT756672** | **MT753209** | **MT739862** | - |
| *Pseudopolydesmus serratus* | MPE01173 | 37.5905 | -76.9638 | Virginia, King William | **MT753984** | - | **MT753214** | **MT739865** | - |
| *Pseudopolydesmus serratus* | MPE01192 | 37.2354 | -78.4649 | Virginia, Prince Edward | **MT753987** | **MT756677** | **MT753217** | **MT739868** | **MT799584** |
| *Pseudopolydesmus serratus* | MPE01427 | 40.8121 | -82.0238 | Ohio, Wayne | **MT753997** | **MT756688** | **MT753230** | **MT739881** | - |
| *Pseudopolydesmus serratus* | MPE01588 | 35.9856 | -79.0238 | North Carolina, Orange | **MT754001** | - | **MT753236** | **MT739888** | - |
| *Pseudopolydesmus serratus* | MPE01646 | 36.9477 | -81.8242 | Virginia, Russell | **MT754002** | - | **MT753237** | **MT739889** | - |
| *Pseudopolydesmus serratus* | MPE01647 | 36.9477 | -81.8242 | Virginia, Russell | **MT754003** | **MT756693** | **MT753238** | **MT739890** | - |
| *Pseudopolydesmus serratus* | MPE01672 | 37.4647 | -81.0623 | West Virginia, Mercer | **MT754005** | **MT756695** | **MT753240** | - | - |
| *Pseudopolydesmus serratus* | MPE01677 | 36.9477 | -81.8242 | Virginia, Russell | **MT754006** | **MT756696** | **MT753241** | **MT739892** | - |
| *Pseudopolydesmus serratus* | MPE01689 | 39.4913 | -82.5797 | Ohio, Hocking | **MT754009** | **MT756699** | **MT753244** | **MT739895** | **MT799590** |
| *Pseudopolydesmus serratus* | MPE01749 | 37.2975 | -82.2987 | Virginia, Dickenson | **MT754011** | **MT756700** | **MT753246** | - | **MT799592** |
| *Pseudopolydesmus serratus* | MPE01750 | 37.8338 | -81.0735 | West Virginia, Raleigh | **MT754012** | - | **MT753247** | **MT739897** | **MT799593** |
| *Pseudopolydesmus serratus* | MPE01779 | 38.722 | -83.433 | Ohio, Adams | **MT754015** | **MT756703** | **MT753250** | **MT739900** | - |
| *Pseudopolydesmus serratus* | MPE01785 | 39.1924 | -86.5105 | Indiana, Monroe | **MT754016** | **MT756704** | **MT753251** | **MT739901** | - |
| *Pseudopolydesmus serratus* | MPE01854 | 38.5675 | -79.1699 | Virginia, Alleghany | **MT754018** | - | **MT753253** | **MT739903** | - |
| *Pseudopolydesmus serratus* | MPE01860 | 38.5948 | -79.1982 | West Virginia, Pendleton | **MT754019** | - | **MT753254** | **MT739904** | - |
| *Pseudopolydesmus serratus* | MPE01864 | 39.6179 | -79.1476 | Maryland, Garrett | **MT754020** | **MT756706** | **MT753256** | **MT739906** | **MT799595** |
| *Pseudopolydesmus serratus* | MPE01878 | 39.0169 | -77.9632 | Virginia, Clarke | **MT754023** | - | **MT753259** | **MT739909** | **MT799598** |
| *Pseudopolydesmus serratus* | MPE01884 | 38.1474 | -78.7443 | Virginia, Albemarle | **MT754024** | - | **MT753260** | **MT739910** | - |
| *Pseudopolydesmus serratus* | MPE01891 | 38.3089 | -79.3603 | Virginia, Augusta | **MT754026** | - | **MT753262** | **MT739912** | - |
| *Pseudopolydesmus serratus* | MPE01894 | 38.33 | -79.2357 | Virginia, Augusta | **MT754027** | **MT756711** | **MT753264** | **MT739914** | - |
| *Pseudopolydesmus serratus* | MPE01913 | 38.2127 | -79.2964 | Virginia, Augusta | **MT754031** | - | **MT753268** | **MT739918** | - |
| *Pseudopolydesmus serratus* | MPE01985 | 38.7739 | -83.1733 | Ohio, Scioto | **MT754036** | **MT756719** | **MT753273** | **MT739923** | **MT799603** |
| *Pseudopolydesmus serratus* | MPE02120 | 39.6733 | -79.7708 | West Virginia, Preston | **MT754040** | - | **MT753277** | **MT739927** | - |
| *Pseudopolydesmus serratus* | MPE02284 | 38.1802 | -81.8384 | West Virginia, Boone | **MT754044** | **MT756727** | **MT753282** | **MT739932** | **MT799609** |
| *Pseudopolydesmus serratus* | MPE02323 | 38.931 | -78.322 | Virginia, Shenandoah | **MT754046** | **MT756729** | **MT753284** | **MT739934** | **MT799611** |
| *Pseudopolydesmus serratus* | MPE02380 | 38.0004 | -78.3703 | Virginia, Albemarle | **MT754047** | **MT756730** | **MT753285** | **MT739935** | **MT799612** |
| *Pseudopolydesmus serratus* | MPE02402 | 34.2393 | -77.8228 | North Carolina, New Hanover | **MT754049** | **MT756734** | **MT753289** | **MT739939** | **MT799614** |
| *Pseudopolydesmus serratus* | MPE02406 | 37.8338 | -81.0735 | West Virginia, Raleigh | **MT754050** | - | **MT753290** | **MT739940** | - |
| *Pseudopolydesmus serratus* | MPE02442 | 37.6801 | -77.9332 | Virginia, Powhatan | **MT754054** | - | **MT753294** | **MT739944** | - |
| *Pseudopolydesmus serratus* | MPE02445 | 38.7666 | -77.4104 | Virginia, Fairfax | **MT754055** | **MT756737** | **MT753295** | **MT739945** | - |
| *Pseudopolydesmus serratus* | MPE02465 | 37.9636 | -78.8991 | Virginia, Nelson | **MT754057** | - | **MT753297** | **MT739947** | - |
| *Pseudopolydesmus serratus* | MPE02466 | 38.8609 | -78.3049 | Virginia, Warren | **MT754058** | - | **MT753298** | **MT739948** | - |
| *Pseudopolydesmus serratus* | MPE02474 | 38.985 | -77.99 | Virginia, Fauquier | **MT754059** | **MT756739** | **MT753299** | **MT739949** | - |
| *Pseudopolydesmus serratus* | MPE02534 | 37.4717 | -80.562 | West Virginia, Monroe | **MT754065** | **MT756744** | **MT753305** | **MT739955** | **MT799619** |
| *Pseudopolydesmus serratus* | MPE02587 | 38.8277 | -78.1882 | Virginia, Warren | **MT754066** | **MT756745** | **MT753306** | **MT739956** | - |
| *Pseudopolydesmus serratus* | MPE02600 | 38.3802 | -78.5036 | Virginia, Greene | **MT754067** | **MT756746** | **MT753307** | **MT739957** | - |
| *Pseudopolydesmus serratus* | MPE02676 | 37.4667 | -83.9164 | Kentucky, Jackson | **MT754071** | **MT756749** | **MT753311** | **MT739961** | - |
| *Pseudopolydesmus serratus* | MPE02713 | 32.5438 | -92.1636 | Louisiana, Ouachita | **MT754074** | **MT756750** | **MT753314** | **MT739964** | **MT799623** |
| *Pseudopolydesmus serratus* | MPE02803 | 37.4964 | -89.3572 | Illinois, Union | - | - | **MT753320** | **MT739970** | **MT799626** |
| *Pseudopolydesmus serratus* | MPE02842 | 38.2301 | -80.2353 | West Virginia, Pocahontas | **MT754080** | - | **MT753323** | **MT739973** | - |
| *Pseudopolydesmus serratus* | MPE02846 | 36.8754 | -77.1825 | Virginia, Sussex | **MT754081** | - | **MT753324** | **MT739974** | - |
| *Pseudopolydesmus serratus* | MPE02874 | 37.778 | -79.1923 | Virginia, Amherst | - | **MT756756** | **MT753326** | **MT739976** | **MT799628** |
| *Pseudopolydesmus serratus* | MPE02915 | 38.6964 | -83.2051 | Ohio, Scioto | - | **MT756757** | **MT753327** | **MT739977** | - |
| *Pseudopolydesmus serratus* | MPE02997 | 39.4211 | -81.361 | Ohio, Washington | **MT754085** | - | **MT753330** | **MT739980** | **MT799630** |
| *Pseudopolydesmus serratus* | MPE03015 | 39.5653 | -79.0631 | Maryland, Garrett | **MT754088** | **MT756760** | **MT753333** | **MT739983** | - |
| *Pseudopolydesmus serratus* | MPE03019 | 39.9079 | -77.4777 | Pennsylvania, Franklin | **MT754089** | **MT756761** | **MT753334** | **MT739984** | - |
| *Pseudopolydesmus serratus* | MPE03020 | 40.0039 | -77.3598 | Pennsylvania, Cumberland | **MT754090** | **MT756762** | **MT753335** | **MT739985** | - |
| *Pseudopolydesmus serratus* | MPE03030 | 41.7093 | -80.8732 | Ohio, Ashtabula | **MT754092** | **MT756764** | **MT753336** | **MT739987** | - |
| *Pseudopolydesmus serratus* | MPE03034 | 42.1774 | -75.4228 | New York, Broome | **MT754094** | **MT756766** | **MT753338** | **MT739989** | **MT799632** |
| *Pseudopolydesmus serratus* | MPE03064 | 40.7805 | -76.1308 | Pennsylvania, Schuylkill | **MT754096** | - | **MT753340** | **MT739991** | **MT799634** |
| *Pseudopolydesmus serratus* | MPE03069 | 41.1034 | -77.2435 | Pennsylvania, Clinton | **MT754098** | **MT756769** | **MT753342** | **MT739993** | - |
| *Pseudopolydesmus serratus* | MPE03073 | 38.5948 | -78.6734 | Virginia, Rockingham | **MT754099** | **MT756770** | **MT753343** | **MT739994** | - |
| *Pseudopolydesmus serratus* | MPE03080 | 42.5433 | -76.6018 | New York, Tompkins | **MT754100** | **MT756771** | **MT753344** | **MT739995** | **MT799635** |
| *Pseudopolydesmus serratus* | MPE03105 | 37.7748 | -83.6825 | Kentucky, Powell | **MT754101** | **MT756772** | **MT753345** | **MT739996** | - |
| *Pseudopolydesmus serratus* | MPE03132 | 38.3047 | -82.3512 | West Virginia, Wayne | **MT754102** | **MT756773** | **MT753346** | **MT739997** | **MT799636** |
| *Pseudopolydesmus serratus* | MPE03138 | 38.3475 | -82.6866 | Kentucky, Boyd | **MT754103** | **MT756774** | **MT753347** | **MT739998** | **MT799637** |
| *Pseudopolydesmus serratus* | MPE03222 | 38.0455 | -83.5494 | Kentucky, Menifee | **MT754105** | - | **MT753351** | **MT740002** | - |
| *Pseudopolydesmus serratus* | MPE03255 | 37.1949 | -85.9411 | Kentucky, Hart | **MT754107** | **MT756777** | - | **MT740004** | - |
| *Pseudopolydesmus serratus* | MPE03308 | 32.3085 | -86.3062 | Alabama, Montgomery | **MT754108** | **MT756778** | **MT753352** | **MT740005** | - |
| *Pseudopolydesmus serratus* | MPE03336 | 39.6793 | -78.7746 | Maryland, Allegany | **MT754112** | **MT756781** | **MT753356** | **MT740009** | - |
| *Pseudopolydesmus serratus* | MPE03439 | 36.467 | -78.429 | Virginia, Madison | - | - | **MT753357** | **MT740010** | - |
| *Pseudopolydesmus serratus* | MPE03443 | 36.55 | -79.35 | Virginia, Pittsylvania | **MT754114** | - | **MT753359** | **MT740012** | - |
| *Pseudopolydesmus serratus* | MPE03522 | 36.6062 | -80.772 | Virginia, Carroll | **MT754118** | - | **MT753364** | **MT740015** | **MT799646** |
| *Pseudopolydesmus serratus* | MPE03534 | 35.8854 | -78.76 | North Carolina, Wake | **MT754119** | - | **MT753365** | **MT740016** | **MT799647** |
| *Pseudopolydesmus serratus* | MPE03546 | 36.2478 | -77.8857 | North Carolina, Halifax | - | - | **MT753368** | **MT740019** | - |
| *Pseudopolydesmus serratus* | MPE03642 | 38.0658 | -77.3803 | Virginia, Caroline | **MT754129** | - | **MT753377** | **MT740028** | - |
| *Pseudopolydesmus serratus* | MPE03867 | 37.6581 | -84.6671 | Kentucky, Garrard | **MT754136** | **MT756800** | **MT753384** | **MT740035** | - |
| *Pseudopolydesmus serratus* | MPE03880 | 38.3788 | -75.7434 | Maryland, Wicomico | **MT754138** | **MT756802** | **MT753386** | **MT740037** | - |
| *Pseudopolydesmus serratus* | MPE03881 | 39.0778 | -76.7712 | Maryland, Anne Arundel | **MT754139** | **MT756803** | **MT753387** | **MT740038** | - |
| *Pseudopolydesmus serratus* | MPE03887 | 38.1359 | -75.4877 | Maryland, Worcester | **MT754140** | - | **MT753388** | **MT740039** | - |
| *Pseudopolydesmus serratus* | MPE03894 | 39.181 | -78.6729 | West Virginia, Hampshire | **MT754141** | - | **MT753389** | **MT740040** | - |
| *Pseudopolydesmus serratus* | MPE03913 | 40.9288 | -91.6694 | Iowa, Henry | - | - | - | **MT740043** | - |
| *Pseudopolydesmus serratus* | MPE03916 | 39.15 | -86.2327 | Indiana, Brown | **MT754144** | - | - | **MT740044** | - |
| *Pseudopolydesmus serratus* | MPE03920 | 40.2095 | -87.648 | Illinois, Vermilion | **MT754145** | **MT756807** | **MT753393** | **MT740046** | - |
| *Pseudopolydesmus serratus* | MPE04030 | 38.2498 | -80.4455 | West Virginia, Greenbrier | **MT754148** | - | **MT753396** | **MT740049** | - |
| *Pseudopolydesmus serratus* | MPE04277 | 30.399 | -91.128 | Louisiana, East Baton Rouge | **MT754155** | - | **MT753405** | **MT740058** | **MT799655** |
| *Pseudopolydesmus serratus* | MPE04609 | 39.615 | -80.9369 | Ohio, Monroe | **MT754166** | **MT756827** | **MT753419** | **MT740072** | - |
| *Pseudopolydesmus serratus* | MPE04667 | 34.7824 | -77.4437 | North Carolina, Onslow | **MT754168** | - | **MT753423** | **MT740076** | - |
| *Pseudopolydesmus serratus* | MPE04711 | 34.7273 | -77.4835 | North Carolina, Onslow | **MT754170** | - | **MT753425** | **MT740078** | **MT799658** |
| *Schizoturanius clavatipes* | MPE04782 | 52.1291 | 86.0769 | Russia, Altai | **MT754172** | - | **MT753431** | **MN232241** | - |
| *Schizoturanius tabescens* | MPE04781 | 50.9831 | 84.2902 | Russia, Altai | - | - | **MT753430** | **MN232242** | - |
| *Scytonotus granulatus* | MPE00979 | 37.3539 | -80.5977 | Virginia, Giles | - | **MT756664** | **MT753197** | **MT739852** | - |
| *Uniramidesmus septimus* | MPE03452 | 44.1491 | 145.8752 | Russia, Sakhalin Oblast | - | - | - | **MN232233** | - |
| **Pyrgodesmidae**  *Myrmecodesmus* sp. | MPE04650 | 18.5841 | -95.0771 | Mexico, Veracruz | - | - | **MT753421** | **MT740074** | - |
